# On the importance of data transformation for data integration in single-cell RNA sequencing analysis

**DOI:** 10.1101/2022.07.19.500522

**Authors:** Youngjun Park, Anne-Christin Hauschild

## Abstract

Recent advances in single-cell RNA (scRNA) sequencing have opened a multitude of possibilities to study tissues down to the level of cellular populations. Subsequently, this enabled various scRNA studies that reported novel or previously undetected subpopulations and their functions by integrating multiple datasets. However, the heterogeneity in single-cell sequencing data makes it unfeasible to adequately integrate multiple datasets generated from different studies. This heterogeneity originates from various sources of noise due to technological limitations. Thus, particular procedures are required to adjust such effects prior to further integrative analysis. Over the last years, numerous single-cell data analysis workflows have been introduced, implementing various read-count transformation methods for de-noising and batch correction. A detailed review of recent single-cell studies shows while many analysis procedures employ various preprocessing steps, they often neglect the importance of a well-chosen and optimized data transformation. This fact is particularly alarming since these data transformations can alter data distribution and thus have a crucial impact on subsequent downstream cell clustering results. Therefore, this study investigates the effects of the various data transformation methods on three different public data scenarios and evaluates them with the most commonly used dimensionality reduction and clustering analysis. Additionally, we discuss its implications for the subsequent application of different deep neural network approaches, such as auto encoders and transfer learning. In summary, our benchmark analysis shows that a large portion of batch effects and noise can be mitigated by simple but well-chosen data transformation methods. We conclude that such optimized preprocessing is crucial and should be the baseline for all comparative single-cell sequencing studies, particularely for integrative analysis of multiple data sets.

## 1 Introduction

Single-cell RNA sequencing (scRNA-seq) enables a high-resolution view of tissues and organisms. With scRNA-seq, it is now possible to understand heterogeneous cell populations by directly sequencing their transcriptome. However, at the same time, observation on a single-cell level results in a higher noise rate in the data due to technological limitations [1]. Although various single-cell technologies are being developed, it remains impossible to capture all the existing RNAs in the cells, since a large proportion of reads are lost during the sequencing preparation steps. Thus, proper post-processing of sequencing data is indispensable [2, 3] and various scRNA-seq data analysis tools were developed and introduced during the last decade. Key steps in the single-cell analysis are composed of the following steps: (1) preprocessing of read-count data, (2) filtering highly variable genes, (3) applying dimensionality reduction (or feature extraction), and (4) clustering on the latent space [4, 5]. All of the available tools have strengths among the above steps. For example, some tools cover the entire process of the above analysis other tools focus on the feature extraction step. Subsequently, well-established dimensionality reduction methods such as t-SNE[6] or UMAP[7] are applied to the feature vectors.

Additionally, following the recent development of deep neural networks (DNN) in computer science, many DNN models were introduced in the area of bioinformatics and single-cell analysis. In particular, autoencoder-based models have been introduced for single-cell RNA sequencing analysis in the last couple of years. These were used as feature encoders to reduce or filter highly variable genes and represent the data with a relatively small size of latent vectors [8]. Thus, these models represent an alternative for feature extraction from high-dimensional data in addition to classical statistical models using singular vector decomposition [9, 10]. Moreover, generative adversarial networks (GANs) based models were developed for single-cell data imputations [11, 12] and data augmentation/generation [13], for instance.

Recently, a number of benchmark studies introduced a variety of different single-cell analysis tools and evaluated their performance, each focusing on specific steps and challenges in single-cell analysis. For instance, Tran *et al*. compared 14 different methods for batch-effect correction. Subsequently, they utilized t-SNE and UMAP for the visualization and quality evaluation of the batch correction [14]. According to their evaluation, the top-three methods for batch mixing were LIGER [15], Harmony [16], and Seurat v3 [17]. Lytal *et al*. compared seven different normalization methods for single-cell RNA sequencing data and evaluated these by k-nearest-neighbor cell type classification. They found that the best performing tools vary between the datasets while Linnorm [18] and scran [19] showed consistent results [20]. Li *et al*. compared four widely used Scanpy-based batch correction methods, and found ComBat [21] performed the best according to their criteria [22]. Luecken *et al*. compared 68 methods and preprocessing combinations using a single-cell dataset including 85 different batches. Their evaluation indicates that, for simple tasks, Harmony [16] is the best choice, and, in more complex tasks, Scanorama [23], and scVI [24] are recommended. Furthermore, they found that deep learning-based models show highly variable performances [25]. Chu *et al*. compared 28 scRNA-seq de-noising methods in 55 scenarios. They developed a set of pipelines for single-cell processing and compared these for various analysis purposes. Their comprehensive benchmark gives the user a practical view of choices [26].

This review of recent benchmark studies highlights the importance of the discussed analysis methods in various tasks. However, the majority of studies neglected the importance of a well-reasoned or optimized choice of appropriate preprocessing methods. The preprocessing step for scRNA read counts usually comprise various data transformation approaches before the data is fed into a complex model for the analysis [4]. For example, Luecken *et al*. solely used scran [19] for normalization and log-transformation for their benchmark dataset. However, although this preprocessing step is considered the best choice for single-cell RNA sequencing analysis, up to now, there is no strong evidence that supports the assumption of generalizability across various datasets, and purposes [3]. Moreover, Cole *et al*. previously reported that there is no one-fits-all solution for every type of single-cell data and pointed out the potential of normalization and data transformation methods for de-noising scRNA-seq datasets [27]. Tian et al. also investigated various analysis pipelines by combining different normalization methods. They also reported that there is not a single best analysis pipeline for all analysis scenarios [28].

Data transformation methods are highly potent tools to alter data distributions and thus have a crucial impact on subsequent dimensionality reduction and clustering analysis [29, 30]. We hypothesize that simple preprocessing steps can reduce batch effects significantly, and each dataset requires different optimal preprocessing. Therefore, comparing evaluation results of different datasets and methods without prior optimization and standardization of preprocessing methods would lead to incomparable outcomes and an unreliable and unfair comparison. Previously, Wang *et al*. compared data transformation methods, log, raw, and z-score, and two different analysis tools, ‘sctransform’ and ‘sc3’. In their result, single-cell clustering analysis results were highly dependent on data transformation [31]. However, their work is limited to a few transformation statistics and methods. Furthermore, they solely focused on the single dataset analysis; hence batch effects or other noise is not considered. Therefore, in the presented study, we aim to fill this gap and investigate the impact of data transformation methods on single and multiple integrated scRNA-seq data analysis tasks, including their benefits for batch effect correction with dimensionality reduction, clustering methods, and their combination with recent DNN models.

## 2 Materials and methods

We tested different combinations of data transformation methods on three different batch effect tasks; (1) single-dataset analysis, (2) multiple datasets analysis, (3) multiple dataset analysis with deep neural networks models, see Figure 2.

**Figure 1:**
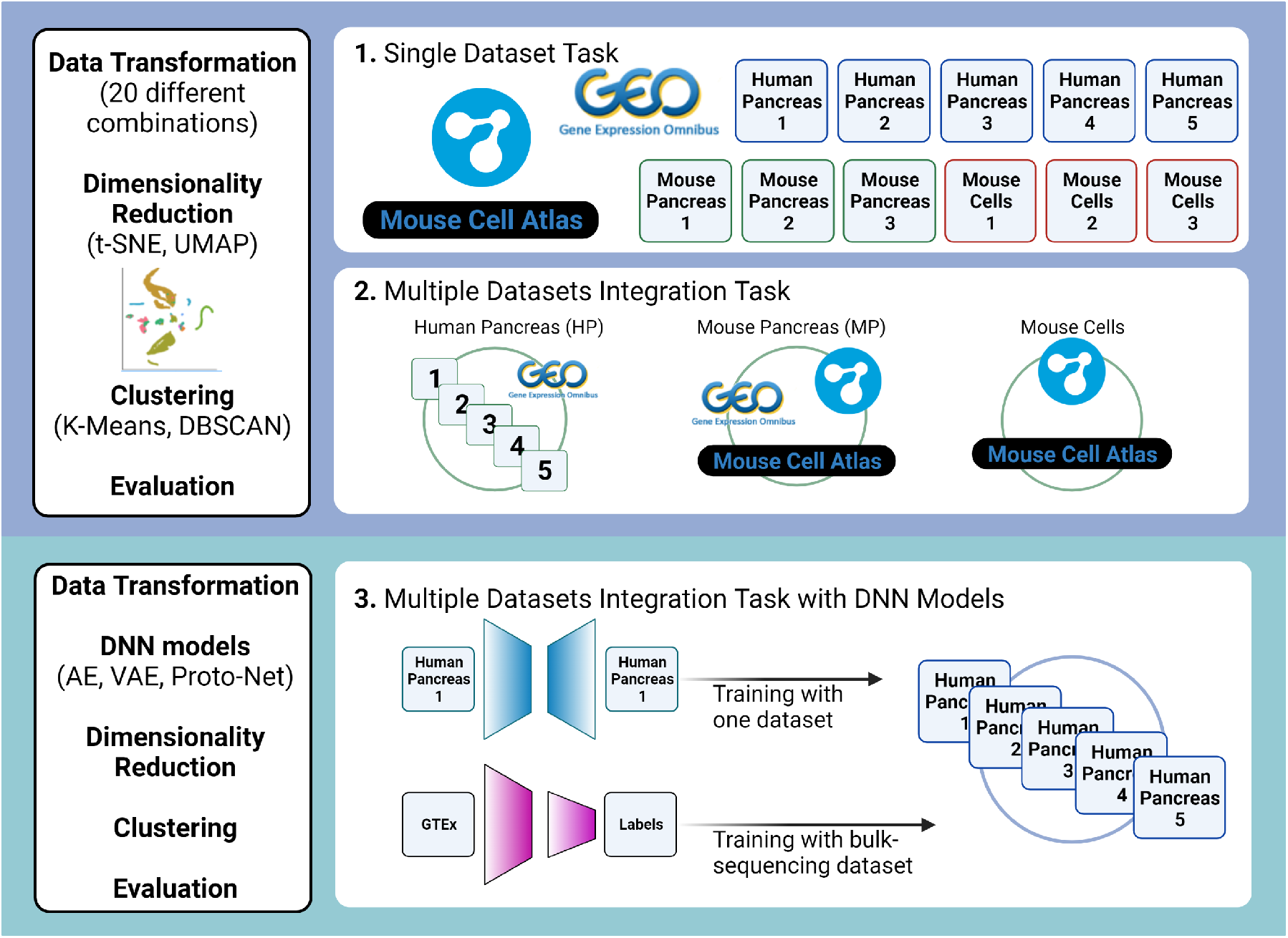
Overview of three main tasks. We tested 20 different combinations of data transformation with two dimensionality reduction methods and two clustering algorithms. In the first task, we evaluated data transformations on a single dataset. On the second task, we integrated datasets and evaluated the batch effect with data transformation methods. On the third task, we used five DNN models on the data integration task to evaluate feature extraction and batch effect correction. Created with BioRender.com.

**Figure 2:**
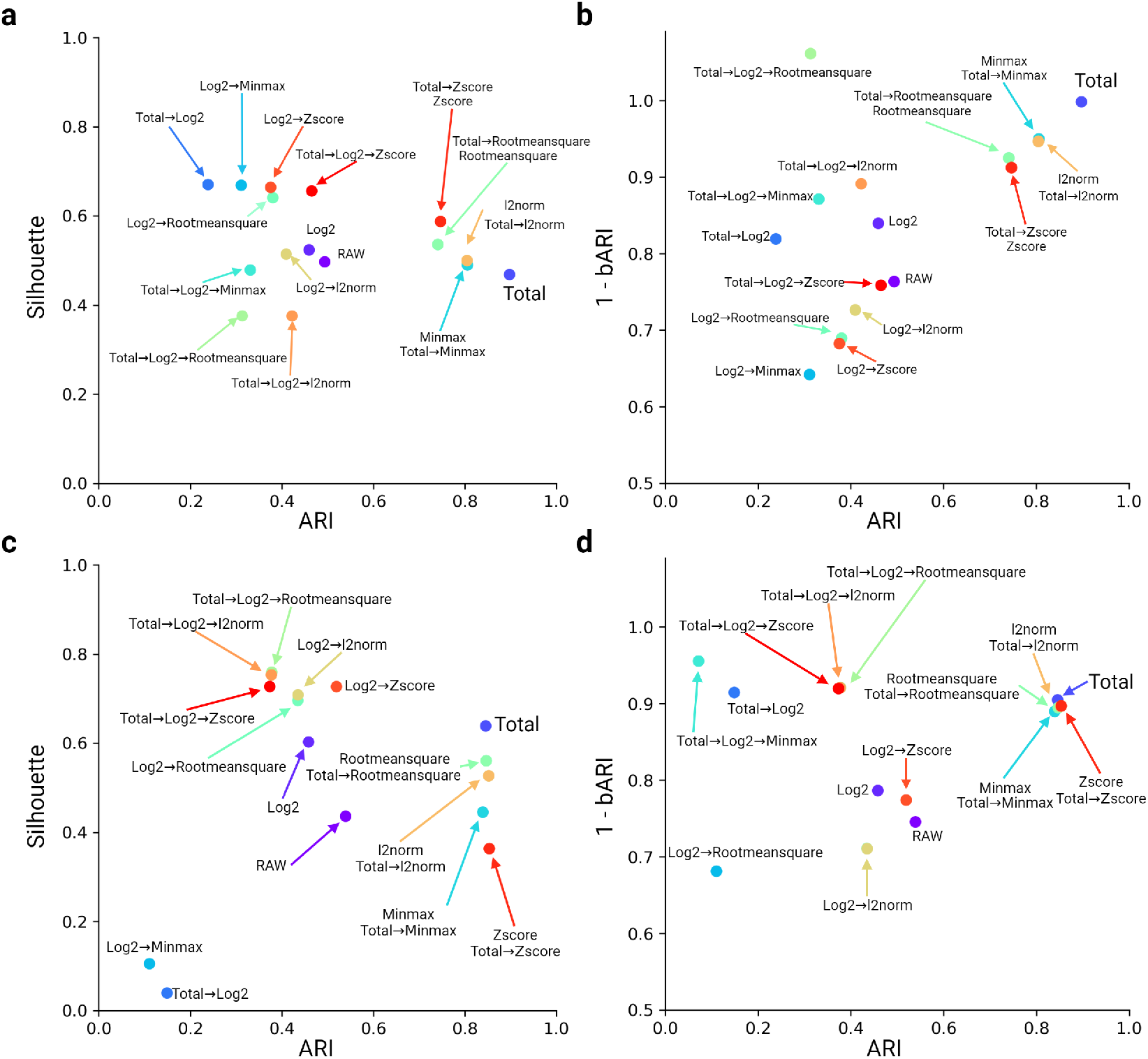
Comparing the impact of 20 different data transformation methods on human and mouse pancreas datasets. **a. b**. The human pancreas dataset consisted of five different single-cell RNA sequencing datasets. Exemplary we show the result obtained using UMAP and DBSCAN clustering method. The full information and visualizations can be found in Supplementary Table S4 and Figure S1. **c. d**. The mouse pancreas dataset consisted of the Tabula Muris and Baron (Mouse) datasets. This shows an example result obtained using t-SNE and DBSCAN. Visualisation result is in the Supplementary Figure S2. All ARI scores are calculated with cell-type labels and clustering results. The bARI scores are calculated based on batch id and clustering results.

### Task 1: Single Dataset Task

During the data transformation benchmark on each data set, whole genes and all cell types were used. First, we analyzed the impact of data transformation on individual datasets. Subsequently, we evaluate the performance of the preprocessing using four analysis pipelines as described in Section 2.3.

### Task 2: Multiple Dataset Integration Task

Multiple datasets analysis was performed on three different subtasks, (1) Human Pancreas Dataset, (2) Mouse Pancreas Dataset, and (3) Mouse Cell Dataset. The aim is to evaluate how much batch effect could be adjusted with simple data transformation methods. Therefore, we will integrate each of the three datasets and employ 20 different data transformation combinations, respectively. Finally, the performance of the preprocessing for all subtasks is evaluated using four analysis pipelines as described in Section 2.3. The batch-ARI score is calculated by considering the biological heterogeneity in addition to technological heterogeneity, and the dimensionality reduction results are also visualized on scatter plots with different colors for different mouse strains.

### Task 3: Multiple Dataset Integration Task with DNN Models

Next, we set a baseline of the Human Pancreas Dataset from the previous analysis. We evaluated the power of various deep neural network models with the same Human Pancreas Dataset. The Autoencoder, Variational Autoencoder [32], and ProtoTypical Network [33] were tested. Lastly, we investigated the impact of transfer learning using a deep neural networks model (Result 3.3).

### 2.1 Dataset description

#### Human Pancreas Dataset

We used five public human pancreas datasets (GSE84133, GSE85241, E-MTAB-5061, GSE81608, and GSE83139). These single-cell RNA sequencing datasets and matching annotation information were downloaded. The datasets are available in varying formats, i.e. GSE84133, GSE83139, and E-MTAB-5061 datasets comprise count data, GSE85241 has adjusted count-like data, and only GSE91608 is provided with normalized RPKM. We summarized the details on the human pancreas datasets in Supplementary Table S1. The download scripts are available in the Github repository, and these are based on Hemberg-lab’s work (https://github.com/hemberg-lab/scRNA.seq.datasets). Before analysis, we exclude unclear cell populations in the dataset from the original study, e.g. ‘unclassified cell’, ‘not applicable’, ‘dropped’, or ‘no label’. For tasks two and three we integrated all five datasets or batches for batch effect analysis. After integration, the integrated dataset comprises 14,918 cells and 15,628 genes. During the batch effect analysis, we used 13 cell types presented in the Baron dataset for the right comparison with recent benchmark work [34].

#### Mouse Pancreas Dataset

We used three different datasets for mouse pancreas cells. This mouse pancreas dataset is composed of the Baron Mouse (inDrop) [35], Pancreas from Tabula Muris’s FACS dataset (SMART-Seq2) [36], and Pancreas from Mouse Cell Atlas dataset (microwell-seq) [37]. The baron mouse and Mouse Cell Atlas datasets and matching annotation information were downloaded via GEO (GSE84133, GSE108097). The Tabula Muris dataset is downloaded from their data portal (https://tabula-muris.ds.czbiohub.org/). Cell type and cluster id information for Mouse Cell Atlas is available on the published supplementary file downloaded from https://ndownloader.figshare.com/files/10760158?private_link=865e694ad06d5857db4b. Each dataset was treated as a different batch. Moreover, we also considered different mouse strains in each dataset. In this case, each mouse strain was treated as a different batch.

For task one, we investigated the batch effect between two mouse strains in the Baron Mouse dataset to compare our approach to a recent benchmark [34]. For task two, we integrated Baron Mouse and Tabula Muris. Labels of the Baron Mouse dataset are converted to make a concordant set with other mouse data: ‘activated stellate’ to ‘stellate’ and ‘quiescent stellate’ to ‘stellate’. Labels of pancreas dataset from Tabula Muris are converted into the same label with Baron Mouse: ‘pancreatic A cell’ to ‘alpha’, ‘type B pancreatic cell’ to ‘beta’, ‘pancreatic D cell’ to ‘delta’, ‘pancreatic acinar cell’ to ‘acinar’, ‘pancreatic ductal cell’ to ‘ductal’, and ‘pancreatic stellate cell’ to ‘stellate’. This integration analysis resulted in 3,213 cells and 13,263 genes. The pancreas dataset from Tabula Muris consisted of four mouse strains. Thus, the batch ARI was calculated based on six mouse strain ids (2 from Baron, 4 from Tabula Muris). Lastly, we integrated Baron Mouse, Tabula Muris, and Mouse Cell Atlas. For the integration, we filtered cells with available cell labels. For better comparability, we excluded the following non-pancreas-related cell types in MCA data, ‘Osteoblast’, ‘Myoblast’, ‘Cycling cell’, ‘Smooth muscle cell’, Stromal cell’, and ‘Epithelial cell’. Labels of MCA were also converted to match Baron and TM. As a result, the integration analysis, including the Mouse Cell Atlas datasets, was done with 5,171 cells and 12,584 genes.

#### Mouse Cell Dataset

For the mouse cell dataset, we used the other Tabula Muris datasets except for the pancreas. This TM dataset contains two different single-cell RNA sequencing datasets using different protocols, SMART-Seq2 from FACS-sorted cells and 10x Genomics platform with CellRanger, and each of the datasets contains additional mouse strain information. Thus, each mouse strain was treated as a batch for the analysis. Among 20 different organs in the Tabula Muris dataset, we filtered a set of tissue/organs having data from both sequencing technologies. Namely, Bladder, Kidney, Limbic muscle, Liver, Lung, Mammary Gland, Marrow, Spleen, Thymus, Tongue, and Trachea were selected. We sampled 200 cells from each tissue and applied data transformation methods. Since each pair shared a set of common cell types, the dataset allowed the evaluation of data transformation methods in terms of batch effect correction. For task two, we integrated expression profiles of the two batches of cells. If the entire dataset was too big, we randomly sampled 2000 cells and performed the analysis.

### 2.2 Data transformation used in single-cell data preprocessing

#### Individual statistics for data transformation

We investigated various data transformation methods applied to scRNA-seq data and chose six data transformation methods. These data transformations are applied to the scRNA-seq data to change its distribution to get better training outcomes. Luecken *et al*. classified data transformation methods into two steps, normalization and transformation [3]. Because of technological limitations that some cells capture more reads and some cells are not, column-wise normalization has been widely applied (total). Minmax normalization is a straightforward method when multiple datasets are integrated. The standardization step using Z-score is also a popular approach. The log transformation could reduce data skewness. Details are listed in Table 2, where E is the expression profile of the cell and e is each of the genes measured in scRNA-seq.

#### Combination of data transformation statistics used for the preprocessing benchmark

Some studies chose an arbitrary data transformation method without further reasoning and fed transformed expression profiles to their complex and novel analysis model. At the same time, we found partial consensus on data transformation methods using three steps: Total*→*Log*→*Z-score. In another point of view, the success of Seurat [38] could contribute to the partial consensus in the scRNA-seq data transformation: Total*→*Log*→*Z-score, see Table 1.

**Table 1:** Data transformation methods. We used six statistics to transform single-cell RNA sequencing data. These statistics are widely used in sequencing data analysis studies.

| RAW | No transformation |
| --- | --- |
| Log2 | $E = \text{Log}_2(e)$ |
| Total | $E = \frac{e}{\text{sum}(e)} * 20000$ |
| $l_2$-norm | $E = \frac{e}{\sqrt{\text{sum}(e^2)}}$ |
| Root mean square | $E = \frac{e}{\sqrt{(\text{mean}(e^2))}}$ |
| Minmax | $E = \frac{e - \min(e)}{\max(e) - \min(e)}$ |
| Z-score | $E = \frac{e - \text{mean}(e)}{\text{std}(e)}$ |

where $e$ is a vector of gene expression level in a cell

**Table 2:**
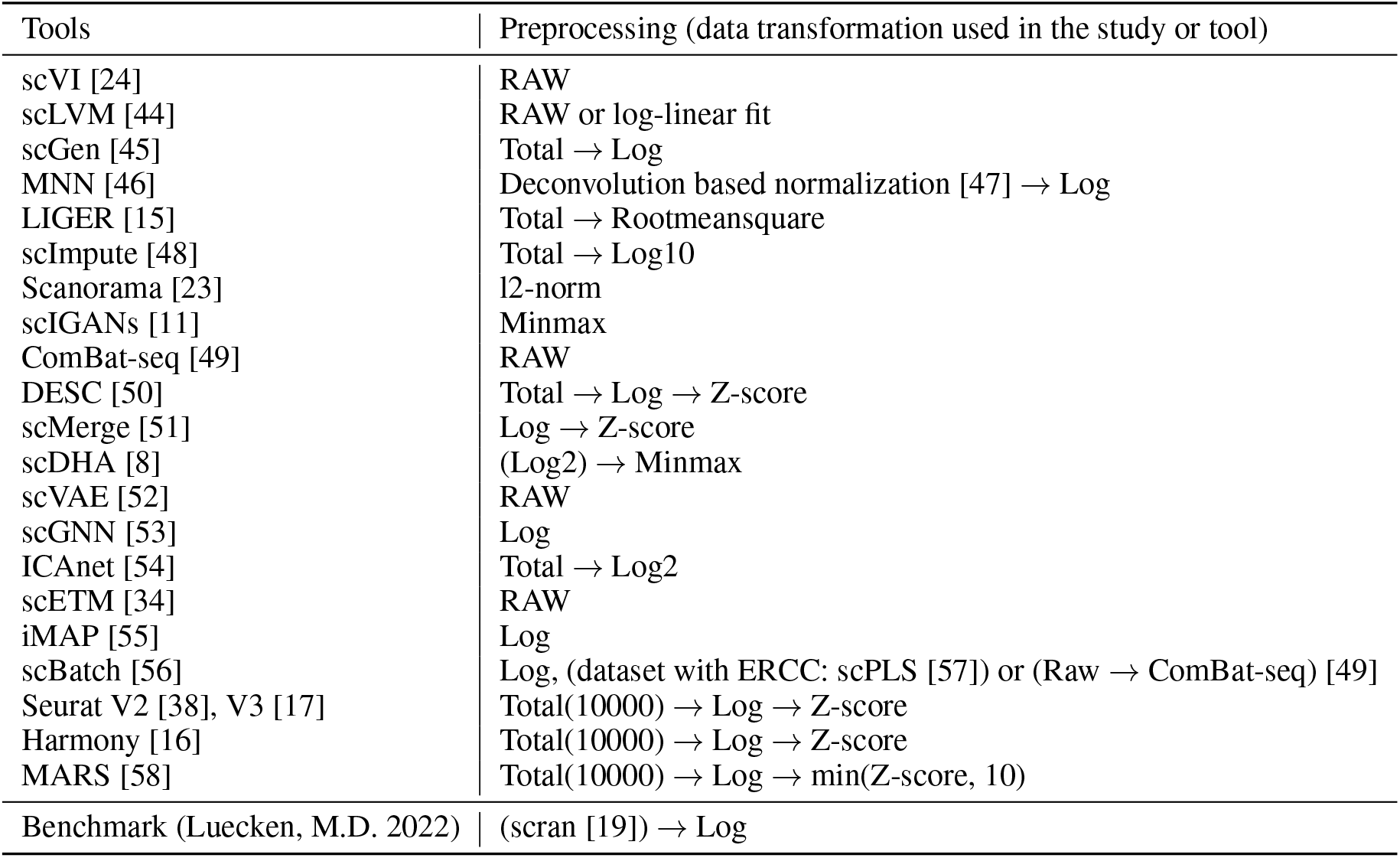
Data transformation methods used in various studies. Summary of recently published studies for single-cell RNA sequencing data and their data transformation methods. Details about statistics is in Table 1. Preprocessing in parenthesis is an optional step depending on dataset. One of the most successful tools for single-cell analysis is Seurat. Thus, many of the studies are using this library [38]. Seurat employs three preprocessing steps: total normalization, log transformation, and z-score standardization.

For the data transformation benchmark performed in this study, 20 different combinations of statistics are employed using six transformation methods: Log2-transformation, Total normalization, Log2*→*Total, Rootmeansquare scaling, Minmax normalization, *l*2-normalization, Z-score transformation. We did not test all possible combinations. But we stick with widely used and plausible data transformation scenarios, normalization, and rescaling.

### 2.3 Single-cell RNA analysis pipelines: dimensionality reduction and clustering

Analysis pipelines are composed of two dimensionality reduction methods and two clustering methods. We used these four analysis pipelines on different tasks with various datasets.

#### Dimensionality Reduction

is a crucial part of single-cell RNA sequencing analysis. High-dimensional gene expression data could be easily projected in 2D space and interpreted. During the last decade, the widely accepted methods are t-SNE[6] and UMAP[7]. Thus, we employed those two methods for our analysis. For t-SNE, we fixed the number of components as two and did not initiate with PCA. For UMAP, we fixed the number of components as two and initiated with the parameter ‘spectral’. For K-Means, we searched for the best ARI score amongst varying parameters for the number of clusters, from half of the number of original labels to the number of original labels + 4. The number of iterations was from 20 to a max of 50. For DBSCAN, we searched for the best ARI score during varying eps parameters from 0.5 to 10 with a 0.5 step size. The parameter for a minimum number of samples is fixed to eight. In all parts of the analysis, we used t-SNE, UMAP, K-Means, and DBSCAN[39] from the python ‘scikit-learn’ packages.

For further analysis, many prior processes are applied to the dimensionality reduction method. This prior process is focused on feature extraction. By doing this, t-SNE or UMAP could effectively represent the cell’s characteristics. The feature selection based on highly variable genes is a widely used prior method [40, 3]. Recently, an autoencoder model was employed for variable gene selection [8]. Furthermore, complex DNN models are introduced for feature extraction. We will cover this DNN model-based feature extraction in the next section (2.4).

#### Clustering Evaluation

Clustering algorithms are often applied to latent space generated from the above dimensionality reduction methods for the identification of cell populations. There are various clustering algorithms. For our analysis, we apply the most widely used K-Means clustering, and DBSCAN [39]. The evaluation of the dimensionality reduction was done with the clustering results using the Adjusted Rand Index (ARI), batch-wise ARI (bARI), and Silhouette score. The ARI is calculated between true cell-type labels and clustering results, and the bARI is calculated between batch ids and clustering results. In the scatter plot, bARI is transformed into 1-bARI and visualized for convenience. The silhouette score is calculated with latent vectors and clustering results. Adjusted Rand Index (ARI), batch-wise ARI (bARI), and Silhouette score are calculated with Python ‘scikit-learn’ packages ‘metrics.adjusted_rand_score’ and ‘metrics.silhouette_score’.

### 2.4 Deep neural networks model

We tested the power of DNN models as a feature extractor for after-dimensionality reduction methods. With this analysis, we incorporate the recent development of DNN model-based tools for single-cell analysis. In this study, we tested five different DNN models, Autoencoder (AE), Variational Autoencoder (VAE), ProtoTypical Network (Proto), Variational Proto (VProto), and TMP-model [41]. We used python and the PyTorch library to implement deep neural network models. The performance of all four types of models was reported on the best model selected from more than 10 training runs using randomly initialized weights. The overall range of performances is shown in the Supplementary Table S2.

#### Autoencoder and Variational Autoencoder

The basic encoder and decoder block is composed fully connected layer, batch normalization layer, and relu layer. In the AE model, there is one hidden layer sized 1024, and the size of the latent layer is tested with 2, 4, 8, 16, 32, 64, 128, and 256. In the case of VAE, the hidden layers have 1024, 128 sized output vectors, and another fully connected layer produces vectors for *µ* and *σ* having 2 to 256 size, similar to AE. The reconstruction loss is calculated with the mean squared error between the original gene expression vector and the reconstructed vector.

#### ProtoTypical and Variational-ProtoTypical Network

ProtoTypical Network is a kind of few-shot learning model having great success in various tasks from computer vision to biomedical analysis [33, 42]. We implemented ProtoTypical Network in two ways, Proto and VProto. The Proto means a general ProtoTypical network that prototypical loss is calculated on latent layer after feature extractor with fully-connected layers. The VProto means that prototypical loss is calculated on the latent layer. Specifically, we obtained a latent vector from the reparametrization trick with *µ* and *σ* from the feature extractor with fully-connected layers and used it to calculate the prototypical loss. The prototypical loss was euclidean distance on latent space.

#### Model Training and Testing Scenario for Batch Effect Analysis

To evaluate the impact of the de-noising power of the data transformation method, we trained the model with only one of five datasets. If the data transformation method has minute power in batch effect adjusting, the model will be easily over-fitted on the training dataset with noise, and evaluation with whole human pancreas datasets would not be good. Furthermore, because the ProtoTypical Network is a supervised learning model [33], utilizing the entire dataset for training and testing simultaneously is nonsense. In this case, we are training the model with one dataset and testing with the other four datasets by comparing cell clusters with the training dataset. For this reason, during the training step in all AE, VAE, Proto, and VProto, we used Baron (Human) dataset and transformed entire human pancreas datasets to visualize and evaluate. With the different sizes of latent vectors, we clustered cells with K-Means and DBSCAN and evaluated their performance in dimensionality reduction and cell population clustering.

#### Transfer Learning Analysis

Recently, transfer learning has been reported as an alternative way to adjust batch effect [41]. The Transferrable Molecular Pattern model (TMP-model) utilized bulk-cell sequencing data and transferred knowledge to the single-cell sequencing task. Its backbone model is another few-shot learning model similar to ProtoTypical network, relation network [43]. We re-evaluated the benefit of transfer learning when datasets become less noisy with the data transformation method. We used the TMP-model pretrained with GTEx (available in https://zenodo.org/record/5529755#.YVGe4bozawF) for the analysis and reproduce previous work (available in https://github.com/iron-lion/tmp_model).

### 2.5 Code and Data Availability

All code and plots used in this analysis are available at the GitHub repository: https://github.com/iron-lion/scRNAseq-preprocessing-impact. Entire datasets are available with SnakeMake file in GitHub/dataset. All visualized results are available in GitHub/notebook. Source codes for the analysis and whole visualization results are available in GitHub/src.

## 3 Results

In order to get an overview of the data transformation methods used for the analysis of single-cell RNA sequencing data, we reviewed 22 recent studies and found a variety of data transformation statistics, see Table 1, in their preprocessing or data cleaning strategies (Table 2).

For example, log transformation is one of the most common data transformation methods in numerous RNA-sequencing data analysis studies. Another widely used method is the min-max normalization, which is especially favored for deep neural network-based models since these computer vision methods use a 0 to 1 range input array. Moreover, the z-score is another method that has been popular since the microarray era [59]. In comparison, column-wise (cell-wise, or total) normalization is the most widely applied method for single-cell analysis due to technological limitations in single-cell sequencing. This limitation makes it impossible to get an evenly distributed read count in each cell. Consequently, each cell has a different number of total read counts. Therefore, total normalization was introduced to handle this issue [60].

Although various data transformations are successfully applied to different studies and tools, there has been no extensive exploration of their impact on the analysis. This gives rise to issues like a lack of comparability or result stability amongst different studies, even on the same dataset. Therefore, we benchmarked 20 different data transformation methods on three tasks with public single-cell datasets, (1) a single dataset analysis task, (2) a multiple datasets integration and batch effect correction task, and (3) a multiple datasets integration task with deep neural networks models 2). Each task was evaluated with regard to the ARI score with its cell-type clustering result on a latent space generated with t-SNE and UMAP (Methods 2.3). For task one, we investigated the impact of the different data transformations on the individual datasets (Result 3.1). Next, we assessed the impact of data transformation on the multiple dataset integration analysis in terms of batch effect correction with three subtasks with popular datasets, Human Pancreas, Mouse Pancreas, and Mouse Cells (Result 3.2) in task two. Finally, for task three, we first set a baseline for the evaluation of the cell-type classification tools and compare it to the performance of deep neural network models in feature extraction and batch effect correction (Result 3.3). Furthermore, the benefit of transfer learning is re-evaluated with data transformation methods (Result 3.3.

### 3.1 Impact of data transformation on the single dataset analysis task

To demonstrate the impact of data preprocessing on the subsequent analysis, we first extended the single dataset analysis task by Wang2020 [31] and Cole2019 [27] with various data transformation methods. In this task, we evaluated 20 data transformations (Methods 2.2) and four single-cell RNA analysis pipelines (Methods 2.3) with eight different datasets.

In the Baron human pancreas dataset, the ARI score showed a high impact of data transformation on the subsequent dimensionality reduction and clustering analysis. Among the 20 different data transformations, Total *→ l*2-norm showed the best ARI with 0.911, and Total *→* Log2 showed the worst with an ARI score of 0.221. The best and worst combinations differ amongst subsequent dimensionality reduction and clustering methods. For instance, for t-SNE+K-Means, *Log*_2_ resulted in the best with ARI 0.808, and Total *→* Log2 was the worst with ARI 0.429. The best results for the analysis with UMAP+DBSCAN, with an ARI 0.904 was achieved by the Total data transformation, and Total*→*Log led to the worst ARI of 0.005. Moreover, for UMAP+K-Means, Rootmeansquare was the best with ARI 0.834, and Total *→* Log2 *→* Minmax was the worst with ARI 0.294. This clearly shows that simple preprocessing methods can tremendously affect single-cell analysis, as demonstrated by an extreme ARI variation ranging between 0.005 and 0.911 on the Baron human pancreas dataset. The overall summary of ARI results is listed in Table 3. The visualization results for each dataset are available online (See Method 2.5).

**Table 3:** The best and worst data transformation method on single scRNA-seq data.

| Analysis | Dataset | Best | ARI | Worst | ARI |
| --- | --- | --- | --- | --- | --- |
| t-SNE<br>K-Means | Baron (Human) | Log2 | 0.808 | Total Log2 | 0.429 |
|  | Muraro | Total Rootmeansquare | 0.921 | Log2 Minmax | 0.569 |
|  | Segerstolpe | Log2 Rootmeansquare | 0.726 | Log2 Minmax | 0.378 |
|  | Wang | Total Log2 Minmax | 0.858 | Log2 Minmax | 0.346 |
|  | Xin | Total Log2 Z-score | 0.722 | Log2 | 0.051 |
|  | Baron (Mouse) | Log2 Z-score | 0.785 | Total Log2 Minmax | 0.163 |
|  | TM (FACS) Pancreas | Total Log2 *l2*-norm | 0.807 | Log2 | 0.461 |
|  | TM (FACS) sampling | Log2 Z-score | 0.610 | RAW | 0.147 |
| t-SNE<br>DBSCAN | Baron (Human) | Total *l2*-norm | 0.911 | Total Log2 | 0.221 |
|  | Muraro | Rootmeansquare | 0.963 | Total Log2 Rootmeansquare | 0.324 |
|  | Segerstolpe | *l2*-norm | 0.950 | Minmax | 0.375 |
|  | Wang | Z-score | 0.872 | Total Log2 | 0.000 |
|  | Xin | Total Minmax | 0.888 | Total Log2 | 0.045 |
|  | Baron (Mouse) | Minmax | 0.929 | Total Log2 | 0.108 |
|  | TM (FACS) Pancreas | Total | 0.774 | Log2 | 0.332 |
|  | TM (FACS) sampling | Total Log2 Z-score | 0.591 | RAW | 0.137 |
| UMAP<br>K-Means | Baron (Human) | Rootmeansquare | 0.834 | Total Log Minmax | 0.294 |
|  | Muraro | Rootmeansquare | 0.957 | Total Log2 Rootmeansquare | 0.246 |
|  | Segerstolpe | Minmax | 0.669 | Log2 Minmax | 0.293 |
|  | Wang | Total Log2 *l2*-norm | 0.856 | Total Log2 | 0.305 |
|  | Xin | Log2 Z-score | 0.638 | Log2 | 0.049 |
|  | Baron (Mouse) | Log2 Meansqaure | 0.669 | Total Log2 | 0.104 |
|  | TM (FACS) Pancreas | Total Log2 Z-score | 0.804 | Log2 Rootmeansquare | 0.403 |
|  | TM (FACS) sampling | Log2 Z-score | 0.637 | RAW | 0.135 |
| UMAP<br>DBSCAN | Baron (Human) | Total | 0.904 | Total Log2 | 0.005 |
|  | Muraro | Total | 0.964 | Log2 Minmax | 0.315 |
|  | Segerstolpe | Total | 0.959 | Total → Log2 Minmax | 0.323 |
|  | Wang | Total Log2 Z-score | 0.863 | Total Log2 | 0.000 |
|  | Xin | Total *l2*-norm | 0.887 | Total Log2 | 0.000 |
|  | Baron (Mouse) | Rootmeansquare | 0.903 | Total Log2 Minmax | 0.000 |
|  | TM (FACS) Pancreas | Total | 0.723 | Log2 | 0.024 |
|  | TM (FACS) sampling | Total Log2 *l2*-norm | 0.519 | RAW | 0.036 |

\* All of visualised results are available online (See Method 2.5)
TM: Tabula Muris dataset / TM sampling: tested with subset of all tissues

The Baron (mouse) dataset contains sequencing data from two different mouse strains. In the recent work [34], this dataset was used to evaluate different strain effect correction tests. It is shown as ‘MP’ in the table for comparison with the benchmark work (Supplementary Table S3). Furthermore, our results show that the noise from different strains can be mitigated when proper data transformation is applied. We obtained an ARI score of 0.929 from Minmax combined with the t-SNE+DBSCAN analysis (Table 3). The Tabula Muris (FACS) dataset contains mouse cell datasets from multiple strains similar to the Baron (mouse) dataset. For all strains, the analysis of the RAW dataset has the worst ARI score. However, when proper data transformation is applied to the TM dataset, the ARI scores improve from nearly

0.147 to above 0.510 in the t-SNE+K-Means analysis, which is consistent with the other analysis pipelines (Table 3). These results also indicate that noise from different batches and mouse strains can be mitigated by the application of proper data transformation.

In summary, the results of our single dataset analysis task confirmed and extended previous findings of Cole *et al*. and Wang *et al*. [27, 31]. There is no method that performs equally well on all datasets. A transformation method that works best for one dataset and a specific analysis pipeline does not necessarily perform well on another.

### 3.2 Impact of data transformation on the multiple dataset integrative task

To reveal the impact of data preprocessing for de-noising and batch effect correction, we evaluated three tasks of popular single-cell datasets with the same 20 data transformations and four single-cell RNA analysis pipelines. The three benchmark tasks are Human Pancreas Dataset, Mouse Pancreas Dataset, and Mouse Cell dataset.

#### 3.2.1 Subtask 1: Human pancreas datasets

The selected human pancreas datasets are frequently used for batch correction tools for single-cell sequencing analysis. They consist of five different single-cell RNA sequencing datasets resulting from four different sequencing protocols (details in Table S1) but are well-labeled with unique cell types in the pancreas. When we investigated the results of all four analysis pipelines, namely t-SNE+K-Means, t-SNE+DBSCAN, UMAP+K-Means, and UMAP+DBSCAN, non of the evaluated data transformation methods worked equally well or best on all. For the method combination of t-SNE and K-Means, Z-score shows the best ARI score with 0.588 and 1-bARI with 0.883. In contrast, for t-SNE and DBSCAN, the Minmax transformation resulted in the best ARI with 0.908 and 1-bARI with 1.019. Moreover, the analysis of UMAP and K-Means demonstrated that Z-score has the best ARI with 0.789 and a 1-bARI of 0.949. Finally, for UMAP and DBSCAN, the Total transformation performed the best with an ARI of 0.898 and 1-bARI of 0.998. All results are available online (See Method 2.5).

Furthermore, we performed a detailed investigation of the UMAP+DBSCAN method combination. When the ARI score is considered, the simple column-wise (or cell-wise) normalization, referred to as Total here, showed the best ARI of 0.898 while RAW or Log2 has mediocre performance (Figure 2a. However, the data transformation method used in Seruat [38], Total *→* Log *→* Z-score, shows better results for the silhouette score than Log2 or RAW but has a worse ARI score of 0.465 (Figure 2a. The bARI score measures how well clustered the dataset is with respect to cell types in contrast to batch. Total *→* Log *→* Rootmeansquare has the best 1-bARI score, but has low ARI score (Figure 2b). The visualization of the results can clearly demonstrate whether the batch effect is mitigated or not. For example, when no transformation is applied (the RAW case), the alpha cell type is scattered into three different clusters, and the beta-cell type is split into four different clusters, with another cell type located between them. However, in the case of Total, the alpha cell type is well clustered, and the beta-cell type is still split but located closely (Supplementary Figure S1.

A recent work by Zhao *et al*. conducted a benchmark analysis using the same dataset [34]. We compared the performance of our result to state-of-the-art methods benchmarked in that work. Notably, the result of our Minmax *→* t-SNE+DBSCAN and Total *→* UMAP+DBSCAN analysis outperformed some of the tools presented by Zhao *et al*. with ARI scores of 0.908 and 0.898 (See Supplementary Table S3 for a detailed comparison).

#### 3.2.2 Subtask 2: Mouse pancreas datasets

For the evaluation of the mouse pancreas task, we used three mouse pancreas data sets, Baron (Mouse) [35], Tabula Muris [36], and Mouse Cell Atlas [37]. At first, we integrated and analyzed the Baron and TM pancreas datasets. For the combined application of t-SNE and DBSCAN, Total data transformation was the best normalization method presenting good batch effect correction performance with an ARI score of 0.865 (Figure 2). In comparison to the human pancreas dataset, there were multiple choices of the data transformation that performed equally well, Rootmeansquare, *l*2-norm, Minmax, and Z-score. Log2 *→* Minmax or Total *→* Log2 methods showed the worst performance. The latent representation of the original data was clearly separated based on the batch labels. However, Total or the other data transformation method above was able to mitigate the batch effect resulting in a good cluster representation (Supplementary Figure S2. In the UMAP and DBSCAN result, Total also showed the best batch effect removal performance with an ARI score of 0.842 and a Silhouette score of 0.639. Similarly, for the combination of t-SNE and DBSCAN, Minmax, Rootmeansquare, *l*2-norm, and Z-score methods showed good performance in batch correction (Supplementary Figure S3).

To further evaluate the impact of preprocessing on the analysis pipelines, we integrated another mouse pancreas dataset from MCA. In particular, we challenged it with an MCA pancreas dataset that does not have a similar set of labels for pancreas-specific cell types. Instead, cell types are aggregated into one label, endocrine cells. Thus, the ARI and bARI score calculated with these labels are not fully comparable with the previous performance of the batch effect correction. Nevertheless, we could observe that the endocrine cells of the MCA dataset are well spread on alpha, beta, and delta clusters (Supplementary Figure S4). Moreover, batch effect correction through data transformation can be observed for the epithelial cell type. In most preprocessing, epithelial cells were not grouped into clear cell clusters. However, in the case of Total*→*Minmax, they are closely located to each other (Supplementary Figure S4.

#### 3.2.3 Subtask 3: Mouse cells datasets

The Tabula Muris dataset consists of two different scRNA-seq datasets, SMART-Seq2 and CellRanger. Therefore, in addition to the technical heterogeneity, there is also biological heterogeneity in each dataset. Thus, we investigated the tissue pairs present in both datasets. Similar to the previous results, the data transformation methods affect the dataset integration of the TM. Depending on the method, resulting cell clusters were either based on the batch label or based on biological cell labels. Significant improvements were observed in 9 out of 11 tissue pairs of the TM dataset (see Figure 3). The results were obtained from the same analysis procedures, DBSCAN clustering on the UMAP. In the case of the cell types bladder, mesenchymal cell (purple) and bladder cell (orange) are represented by two different clusters in ‘RAW’. These clusters are strongly influenced by the batch effect and thus can be labeled by SMART-Seq2 and CellRanger. However, after data transformation with log2 *→* Minmax normalization, the cells clustered well based on the cell labels, and the batch labels were mixed (see Figure 3). The ARI score improved from 0.347 to 0.812, and the bARI score decreased from 0.268 to −0.002. Similarly, the raw dataset showed batch-associated label clusters. However, after transforming the scRNA-seq data with Total *→* Log2 *→ l*2-norm, we were able to identify cell label-associated clusters based on the latent space using UMAP. However, the cell types bone-marrow and tongue showed relatively marginal batch effect correction for all 20 data transformation methods. In the case of bone-marrow, when UMAP and DBSCAN were used, the ARI of bone-marrow improved from 0.211 to 0.316 (Rootmeansquare), and when t-SNE and DBSCAN were used, ARI improved from 0.282 to 0.533 (Log2 *→ l*2-norm). In the case of tongue, UMAP and DBSCAN analysis showed the changes from 0.182 to 0.241 (total *→* Z-score), and t-SNE and DBSCAN showed the changes from 0.180 to 0.283 (*l*2-norm). Full plots for each of the tissues are available online (See Method 2.5).

**Figure 3:**
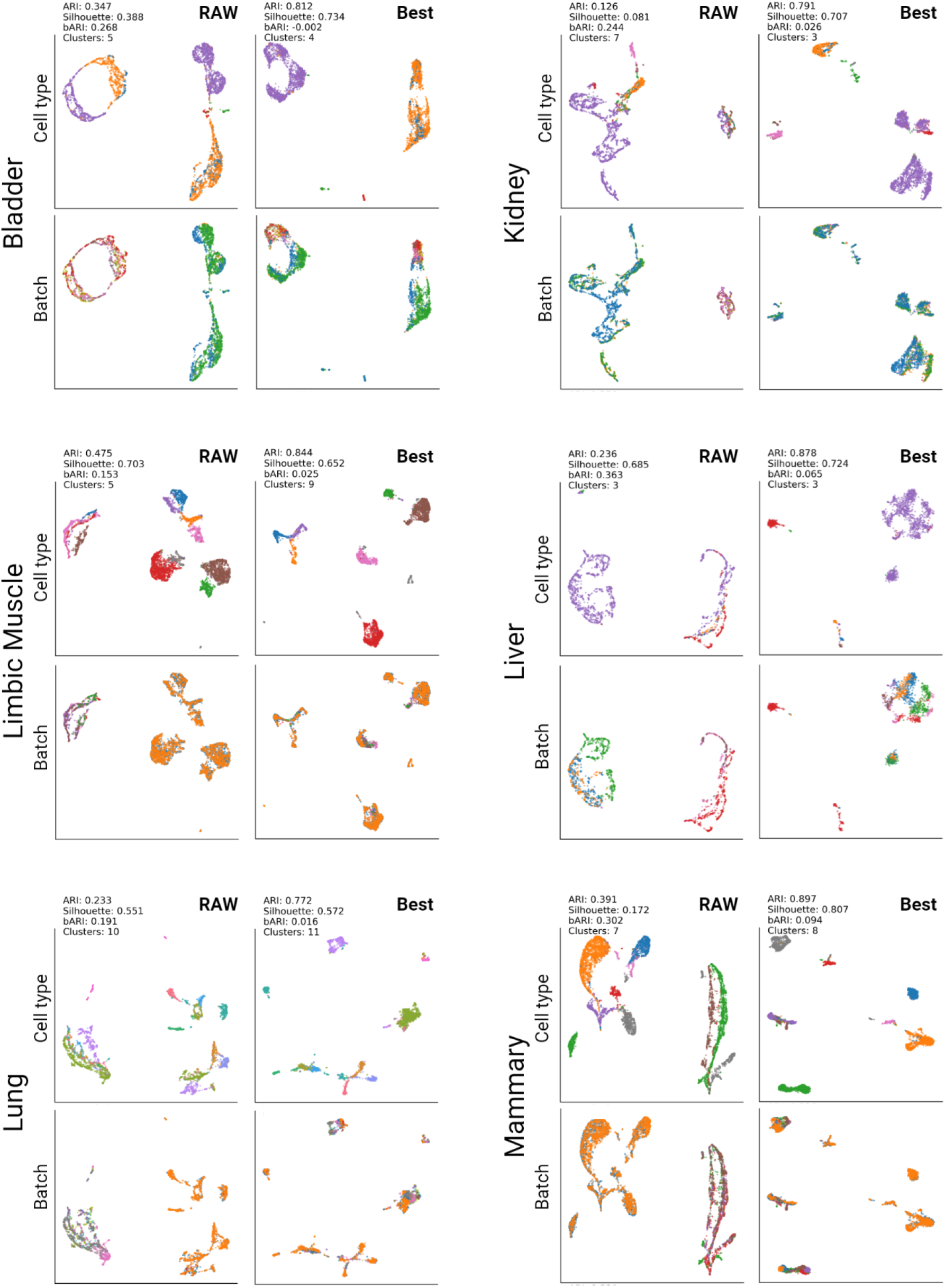
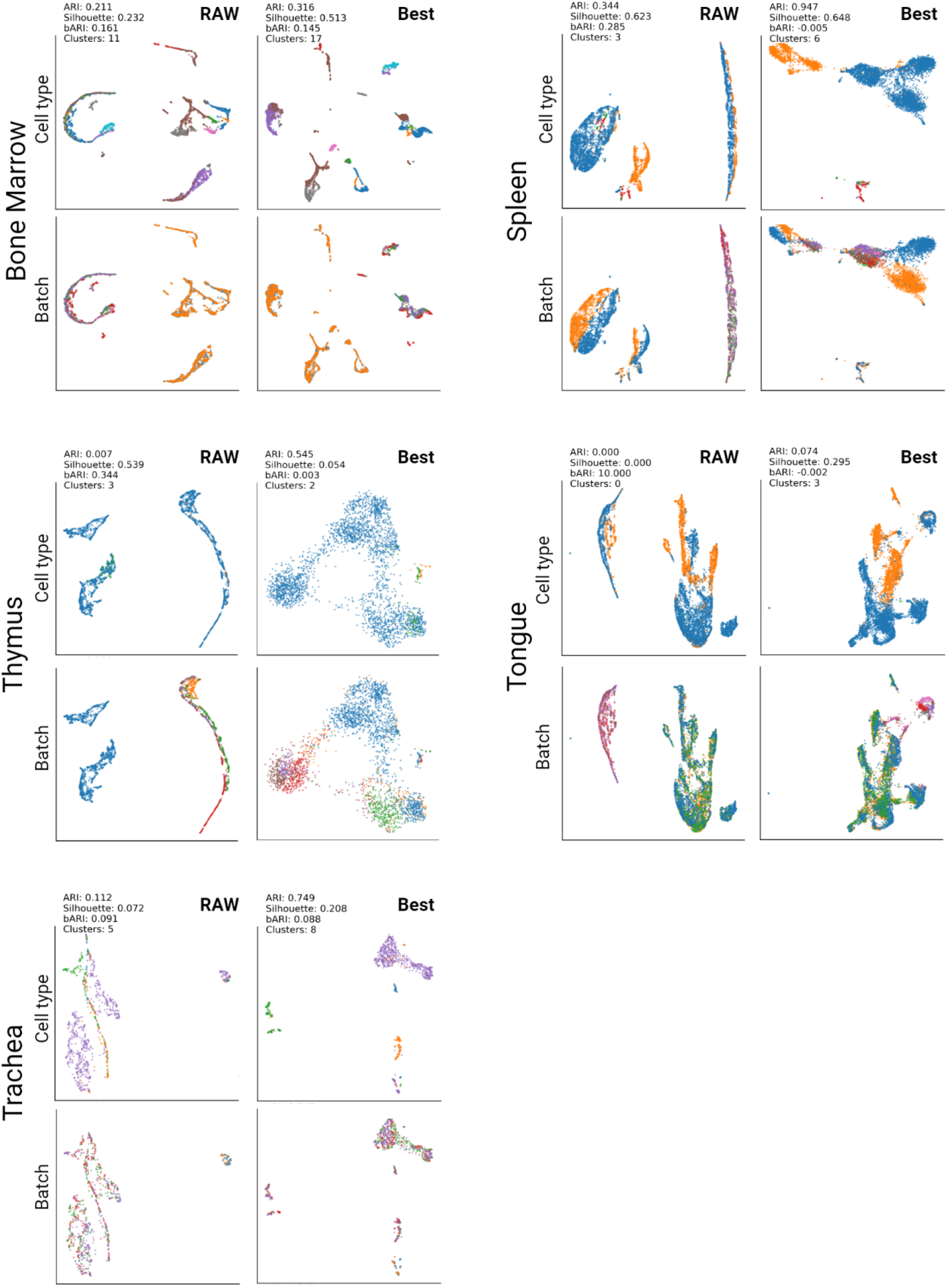
Visualization results for the Tabula Muris dataset. UMAP and DBSCAN was used for this analysis. ‘RAW’ indicates the original read count data of TM. ‘Best’ represents the best result out of 20 data transformation methods. The cell types and batch ids are represented with different colors in each plot. The entire results are shown in Supplementary Table S5 and all plots and legends are available online (See Method 2.5).

In addition to the Tabula Muris dataset, we integrated the Mouse Cell Atlas dataset, which integrates single-cell RNA sequencing datasets from mouse organs and tissues. However, as described in the previous subtask, the Mouse Cell Atlas dataset has a different level of cell labels compared to the other datasets. Thus the three datasets shared only a few common cell labels. Thus the ARI and bARI are not dramatically improved. However, we observed a varying representation of cell clusters in latent space depending on the data transformation method. Therefore, by changing the data transformation method, it was possible to improve the latent space representation and subsequently improve clustering results (Data available online (See Method 2.5).

### 3.3 Impact of data transformation on the DNN model evaluation task

To demonstrate a proper comparison for the model evaluation, we built various neural network models and evaluated their performance by comparing them with the above results. Here we consider the best benchmark result as a baseline for the given human pancreas dataset, we assessed the power of the DNN model.

Previously, various tools using deep neural network-based models have been developed for single-cell analysis. Moreover, autoencoder is another widely used method to extract features, reduce the dimensionality of sequencing data, and represent cell types with feature vectors. Depending on the implementation details, some tools produce a 2-dimensional vector representation like t-SNE and UMAP. Others focus on feature extraction itself and require t-SNE and UMAP as a further dimensionality reduction step for 2-dimensional representation.

We implemented the simplest neural network-based model as a feature extractor. With this DNN model task, we were able to evaluate the importance of data transformation methods in terms of ‘Garbage In, Garbage Out’. At first, we tested an untrained neural network model. The simplest neural network with two fully-connected layers initialized with random weights was built on the RAW as well as the best performing ‘Total’ transformed single-cell RNA-seq data of the human pancreas dataset. In the latent space of the DNN trained on the ‘RAW’ data, the same cell types are segregated amongst different clusters that are based on batch number. This resulted in an ARI score of 0.542 using t-SNE+DBSCAN and 0.420 using UMAP+DBSCAN with latent size 128 and one hidden layer having a size of 1024. In contrast, the latent space of the DNN trained on the ‘Total’ transformed data showed a significantly better clustering of cell types with an ARI score of 0.891 at latent vector size 128. While the randomness of the weight initialization step in every run has produced slightly different numbers, the data transformation step consistently led to a significant improvement of the model performance.

Moreover, to investigate the feature extraction power of autoencoder models, we tested autoencoder (AE) and variational autoencoder (VAE) with two layers for encoder and decoder (see Method section for details). Here, we used only the Baron dataset for training and evaluated the model on the remaining entire HP dataset. Our results show that the DNN model learns to extract features about cell types only within the Baron data. If the batch effect is not mitigated by preprocessing with data transformation, the DNN model is not able to extract proper signatures for cell types in another dataset (e.g. the HP). The performance of the autoencoder without data transformation was 0.625 ARI using t-SNE+DBSCAN and 0.551 ARI using UMAP+DBSCAN (Supplementary Figure S5). The ‘Total’ transformed data showed a better performance than the ‘RAW’ data even in combination with the AE and VAE DNN models. The combination of the ‘Total’ transformed data with an AE model resulted in a best ARI score of 0.947 (Supplementary Figure S6). Similarly, the best result in combination with a VAE showed an ARI of 0.943 with a latent size of 128 with a hidden layer having a size of 1024 (Supplementary Figure S7). In summary, when we compared the result with the previous analysis, using the DNN model could improve the clustering result (RAW 0.494 *→* 0.551 / Total 0.898 *→* 0.947). However, the data transformation has more impact on the clustering results. When we compared our simple DNN model with recent benchmark work, AE and VAE using Total *→* UMAP+DBSCAN showed comparably good performance with ARI 0.938 and 0.942 (Supplementary Table S3).

In the next step, we used the cell labels for the evaluation of supervised dimensionality reduction methods. Therefore, we modified AE and VAE to calculate the ProtoTypical loss. Similar to the previous task, the training was conducted solely using the Baron dataset and its associate cell type labels. With the ProtoTypical loss, the impact of data transformation is also more critical than the model complexity. The results shown in Figure 4 demonstrate that the use of the raw dataset leads to overfitting to the batch noise and subsequent poor cell type clustering (Figure 4, Supplementary Figure S8). We tested five different sizes for the latent vector from 2 to 32 while a hidden layer is fixed with a size of 1024. At the size eight, we could find convergence (Figure 4). While it remains to be evaluated whether more complex encoder/decoder models and more optimized parameters could result in performance improvements, our results indicate that latent space sizes between 8 and 32 are reasonable and 50 or 100 is also enough for the size of latent vector as reported in the other studies [24, 34, 52, 8].

**Figure 4:**
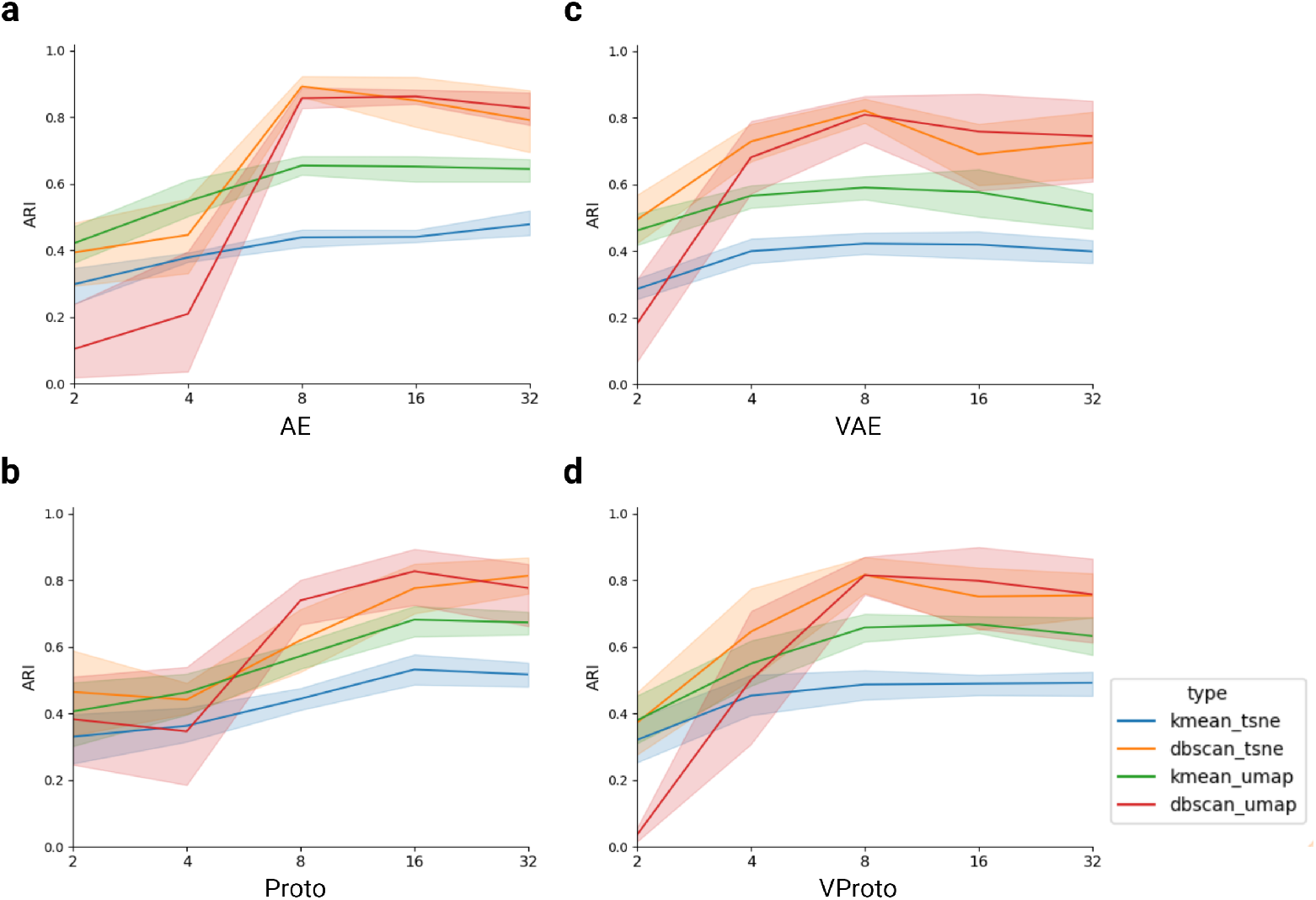
Evaluation of the performance of deep neural network models with varying sizes of the latent vector. Shown ARI scores are obtained after 200 episodes of MAML training with Baron (Human) datasets. All human pancreas datasets are transformed with Total. The x-axis corresponds to the size of the latent vector. The colored area represents a 95% confidence interval with more than 6+ repeats of training and evaluations. Abbreviation: Autoencoder (AE), Variational Autoencoder (VAE), ProtoTypical Network on AE backbone (Proto), ProtoTypical Network on VAE backbone (VProto).

In our previous work [41], we presented transfer learning-based approaches by exploiting bulk sequencing datasets to compensate for an issue from the limited data condition with single-cell RNA sequencing data. The results showed an improvement in the few-shot classification task when utilizing the GTEx dataset as a baseline model for the human pancreas dataset. However, when we applied a different data transformation step prior to the analysis, for some datasets, the significant benefit from the transfer learning did not persist. Similar to the previous analysis, we tested the impact of the different data transformation methods. In concordance with the previous outcomes the accuracy of the cell type classification varies depending on the chosen data transformation (Figure 5a). In this study, The Total showed better baseline performance with the pre-trained model from the GTEx dataset. As a result, when we reproduced fine-tuning step in TMP-model with different data transformations, we could observe that, in some datasets, the performance gained from transfer learning disappeared. This result is evident in the ‘Segerstolpe’ dataset case (Figure 5b).

**Figure 5:**
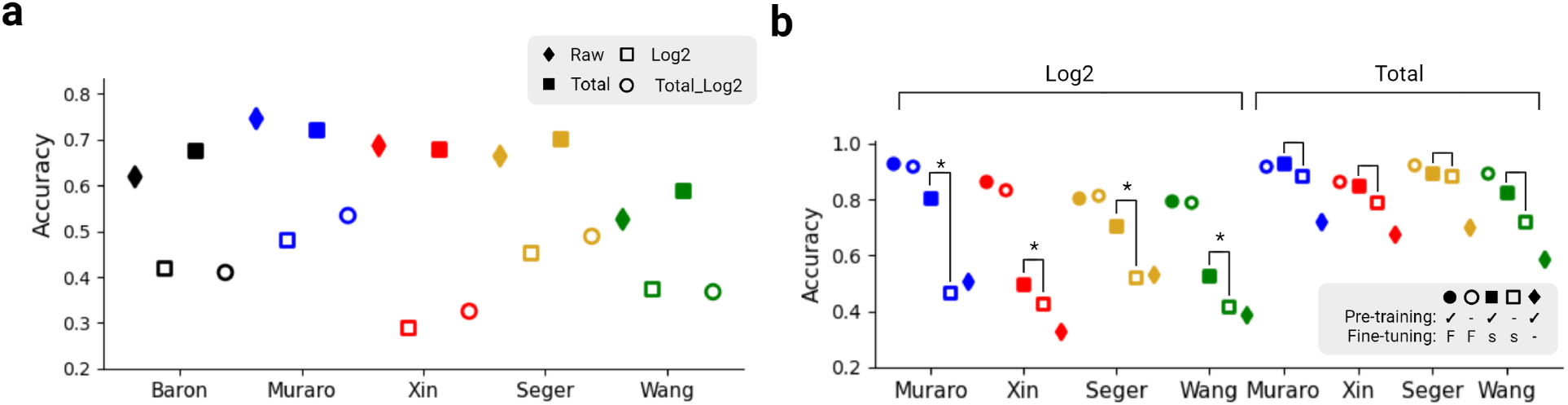
Data transformation impact on evaluation of transfer learning. **a**. Data transformation methods impact on classification task in each of human pancreas data. Details of transformation methods are in Table 1. **b**. TMP-model performance comparison on Log2 and Total methods. Pretraining indicates that model initialized with GTEx-pretrained networks, and Fine-tuning ‘F’ indicates entire dataset of Baron was used on fine-tuning step and ‘s’ indicates only 15 cells for each class was used on fine-tuning step.

## 4 Discussion

Today, single-cell analysis is an accessible and viable option for many research areas and scientific questions. Due to this trend, various single-cell analysis tools emerged. The developed tools typically comprise preprocessing as well as analysis methods. The preprocessing steps can, for example, implement data transformation and de-noising, while analysis steps often include dimensionality reduction and clustering. Not all scRNA-seq analysis tools follow the same structure and order of steps, yet preprocessing and subsequent analysis can be considered as two separate steps with different goals. As we previously described, each tool has slightly different methods not only for data transformation but also for de-noising and batch effect correction, dimensionality reduction, clustering, and classification. Recently, complex DNN models have been applied for analysis that demonstrated better performance than previous probabilistic models [25].

Our review of recent literature showed that even for fairly straightforward tasks like cell-type identification, where often few genes can serve as marker genes, many studies choose different preprocessing and analysis methods without adequate justification. Especially, the data transformation as the very first step of a preprocessing and analysis pipeline was not well investigated in the many benchmark papers for de-noising [14, 25, 20]. Previously, Wang *et al*. reported the impact of preprocessing on single-cell analysis; however, they did not cover an integrative analysis of multiple datasets where batch effects are a tremendous challenge [31]. Because data transformation is usually the first preprocessing step, it can affect the entire analysis process. Our results demonstrate that most of the noise hindering cell-type identification on t-SNE and UMAP space can be easily removed by applying simple but well-chosen data transformation with well-known statistics. Furthermore, according to our observations, cell population representation changed dramatically depending on the analysis pipeline and data transformation method. Our results confirm the findings of Ahlmann-Eltze and Huber, which showed changing marker gene distribution depending on the data transformation [30]. For benchmark works, this indicates that a lack of optimization of the data transformation method within the analysis pipelines can lead to a distorted comparison of results. For instance, the benchmark work by Luecken *et al*. tried to find an optimal preprocessing step for various tools [25]. However, they only set a ‘RAW’ as a baseline of their benchmark work. Our results indicate that it is crucial for future benchmark studies to incorporate an optimization step for proper data transformation in the analysis pipelines to prevent distorted comparisons between various methods. Moreover, this indicates that further detailed evaluations of the robustness and performance of current single-cell RNA sequencing analysis tools are required. To overcome this issue, many studies aimed to adjust the gene-expression profiles to various statistical distributions. However, finding the true underlying distribution of a single cell transcriptome presents to be infeasible unless we fully understand the true distribution of the entire transcriptome in cells with various cell states. Therefore, if a baseline distribution can not be determined, we suggest resorting to heuristics, particularly searching for a proper baseline using an optimized data transformation before proceeding with model development, optimization, evaluation, and comparison.

Meanwhile, our results suggest that t-SNE and UMAP are still powerful tools for single-cell analysis that can compete with more recent approaches. This confirms the results of a recent work by Ahlmann-Eltze and Huber. They found that a simple logarithm with principal component analysis showed well-separated cell clusters in the latent space [30]. Lause *et al*. discussed how data transformation or scaling can affect the gene selection step and its downstream analysis. They pointed out that highly variable gene selection methods usually use their mean and variance value because neither log-transformation nor square-root transformation is sufficient for variable stabilization. In particular, they showed that the analytic Pearson residuals method works best for variable gene selections, but log-transformation also had good performance [61]. However, in our study, we excluded the variable gene selection step to retain a clear view of the impact of data transformation on the t-SNE and UMAP analysis. We demonstrated that without any complex model for feature extraction or feature selection, well-combined data transformation and dimensionality reduction methods show good performance in cell type classification tasks. Thus, in a future analysis, we aim to investigate further the interplay of data transformation quality and feature selection methodology.

Finally, we investigated the potential of DNN-based models for single-cell analysis and compared the findings of tasks one and two to an example for more elaborate analysis methods. The DNN models were able to compress gene expression profiles into very small-sized vectors and made it possible to project efficiently onto a 2D-latent space for visualization and clustering. These new methods can improve the performance of t-SNE and UMAP with their feature selection power. In the HP dataset, the ARI improved from 0.898 (Total) to nearly 0.947 (the best from AE). A recent benchmark study [34], reported an ARI score of 0.968 for the best result using the Seruat[17] approach on the same HP dataset (Supplementary Table S3). However, some of the models proposed in their study show worse ARI scores compared to our straightforward analysis pipeline, namely, Total *→* UMAP+DBSCAN. This suggests that their result might improve if an adequate data transformation step is used and the model is optimized for newly transformed data. Furthermore, we aimed to check the potential of the supervised dimensionality reduction method with a ProtoTypical loss. The ProtoTypical loss allowed us to fully utilize the class label. This supervised dimensionality reduction model can be adjusted for specific research questions. Our findings suggest that in circumstances where the datasets are already well understood, ProtoTypical network models can be a good option to investigate underlying biological meanings. For example, finding novel gene markers for specific cell types. The potential of this kind of approach is also discussed in recent work by Chari et al. or Chen et al. [62, 63].

In the computer vision field, transfer learning has been employed, for example, to transfer knowledge from IMAGENET to medical imaging [64, 65]. However, a recent study indicated benefits in terms of training time-cost for transfer learning but did not show advantages in terms of performance (or accuracy) when all parameters are well optimized, and the dataset size is sufficient [66]. To confirm these findings, we investigated the impact of data transformation in recent transfer learning studies. In a study by Zhao *et al*., they presented a transfer learning approach between mouse and human pancreas datasets [34]. They accomplished an ARI score of 0.868 on the mouse-to-human transfer learning task and an ARI score of 0.800 on the human-to-mouse transfer learning task. The transfer learning result was not better than their unsupervised clustering analysis. The TMP-model of Park *et al*. indicates that transfer learning could adjust batch effects on various datasets [41]. However, a re-investigation employing different data transformation methods for preprocessing revealed that large performance gains can be achieved by a properly chosen optimal data transformation method. Compared to optimal preprocessing, the employed transfer learning models solely show a slight gain in performance, if at all. This suggests that transfer learning might not have significant benefits when preprocessing steps are well optimized. These two observations represent a concordant conclusion with the transfer learning studies conducted on medical imaging [66].

In conclusion, there is a need for sophisticated models to uncover complex hidden knowledge from large-scale datasets. When more and more scRNA-seq datasets accumulate, data-driven analysis becomes the only viable and indispensable option for biomedical research. Our study demonstrates that data preprocessing is a crucial step for integrative data analysis and requires optimization for the subsequent analysis pipeline. We envision that our report will guide future integrative data analysis and also help complex model development by proposing the correct baseline accuracy.

