## Supplementary material for "On the importance of data transformation for data integration in single-cell RNA sequencing analysis": 220719_Benchmark_Supplementary.pdf

### 1 Supplementary Figures and Tables


- [Table S1] Single-cell human pancreas datasets description
- [Figure S1] UMAP representation for comparison of 20 different data transformation
- [Table S2] DNN models' performance range from 10+ repeats with random seed
- [Table S3] Cell-type clustering performance comparison with the other study
- [Table S4] Comparison of 20 different data transformation on human pancreas datasets
- [Figure S2] Mouse Pancreas analysis with t-SNE and DBSCAN
- [Figure S3] Mouse Pancreas analysis with UMAP and DBSCAN
- [Figure S4] Three mouse pancreas datasets analysis
- [Table S5] Entire data of two batches integration analysis with Tabula Muris
- [Figure S8] The performance of deep neural networks models during varying the size of latent vectors
- [Figure S5] Latent representation using Autoencoder without data transformation
- [Figure S6] Latent representation using Autoencoder with Total data transformation
- [Figure S7] Latent representation using Variational Autoencoder with Total data transformation

Table S1: **Single-cell human pancreas datasets description.**

| Dataset | Number of class | Protocol | Accession ID | Ref |
|---|---|---|---|---|
| Baron (Human) | 14 | inDrop | GSE84133 | [35] |
| Muraro | 10 | CEL-Seq2 | GSE85241 | [67] |
| Segerstolpe | 14 | SMART-Seq2 | E-MTAB-5061 | [68] |
| Xin | 8 | Fluidigm C1 | GSE81608 | [69] |
| Wang | 7 | SMART-seq | GSE83139 | [70] |

Table S2: **DNN models' performance range from 10+ repeats with random seed.**

| | Reconstruct | | MAML | |
|---|---|---|---|---|
| | RAW | Total | RAW | Total |
| AE-tSNE-KMean | 0.315∼0.470 | 0.385∼0.588 | 0.315∼0.459 | 0.433∼0.532 |
| AE-tSNE-DBSCAN | 0.402∼0.591 | 0.468∼0.906 | 0.390∼0.615 | 0.606∼0.898 |
| AE-UMAP-KMean | 0.416∼0.521 | 0.550∼0.710 | 0.384∼0.507 | 0.686∼0.733 |
| AE-UMAP-DBSCAN | 0.406∼0.599 | 0.546∼0.870 | 0.464∼0.568 | 0.836∼0.947 |
| VAE-tSNE-KMean | 0.219∼0.346 | 0.432∼0.522 | 0.356∼0.468 | 0.454∼0.571 |
| VAE-tSNE-DBSCAN | 0.194∼0.618 | 0.558∼0.897 | 0.382∼0.578 | 0.520∼0.925 |
| VAE-UMAP-KMean | 0.229∼0.370 | 0.611∼0.785 | 0.419∼0.495 | 0.537∼0.657 |
| VAE-UMAP-DBSCAN | 0.063∼0.387 | 0.444∼0.894 | 0.438∼0.575 | 0.526∼0.943 |
| Proto-tSNE-KMean | NA | NA | 0.359∼0.521 | 0.418∼0.632 |
| Proto-tSNE-DBSCAN | NA | NA | 0.456∼0.574 | 0.711∼0.933 |
| Proto-UMAP-KMean | NA | NA | 0.429∼0.502 | 0.641∼0.737 |
| Proto-UMAP-DBSCAN | NA | NA | 0.476∼0.576 | 0.845∼0.907 |
| VProto-tSNE-KMean | NA | NA | 0.356∼0.488 | 0.454∼0.586 |
| VProto-tSNE-DBSCAN | NA | NA | 0.470∼0.597 | 0.587∼0.933 |
| VProto-UMAP-KMean | NA | NA | 0.484∼0.533 | 0.620∼0.740 |
| VProto-UMAP-DBSCAN | NA | NA | 0.494∼0.587 | 0.669∼0.903 |

Reconstruct: 20 epochs, batch size 64, lr=0.5, latent vector size 128
MAML: 200 episodes, batch/query size 5, lr=0.0005, latent vector size 128
∗ The model was trained with Baron dataset and tested with HP dataset.

Table S3: **Cell-type clustering performance comparison with the recent benchmark study.**

| Tools | MP | HP |
|---|---|---|
| Harmony ∗ | 0.969 | 0.955 |
| Scanorama ∗ | 0.915 | 0.859 |
| Seruat ∗ | 0.944 | 0.968 |
| scVAE-GM ∗ | 0.805 | NA |
| scVI ∗ | 0.932 | 0.759 |
| LIGER ∗ | 0.914 | 0.911 |
| scVI-LD ∗ | 0.875 | 0.656 |
| scETM ∗ | 0.946 | 0.943 |
| scETM $-\lambda$ ∗ | 0.851 | 0.474 |
| scETM + adv ∗ | 0.944 | 0.946 |
| Minmax + t-SNE+ DBSCAN | 0.929 | 0.908 |
| Total + UMAP+ DBSCAN | 0.848 | 0.898 |
| Rootmeansquare + UMAP+ DBSCAN | 0.903 | 0.741 |
| $l2$-norm + UMAP+ DBSCAN | 0.902 | 0.804 |
| Total + AE + UMAP+ DBSCAN○ | NA | 0.947 |
| Total + VAE + UMAP+ DBSCAN ○ | NA | 0.943 |

MP: Baron (Mouse) data
HP: Human pancreas datasets
∗ data are directly derived from the result by Zhao *et al.* [34]
○ is trained with Baron, and picked best result of ten repeats

Table S4: **Comparison of 20 different data transformation on human pancreas datasets.** Five different Human pancreas datasets are integrated and analyzed using dimensionality reduction and clustering methods. Best ARI score amongst different data transformation is marked with green background color.

| | t-SNE & K-Means | | | t-SNE & DBSCAN | | | UMAP & K-Means | | | UMAP & DBSCAN | | |
|---|---|---|---|---|---|---|---|---|---|---|---|---|
| | ARI | Sil. | bARI | ARI | Sil. | bARI | ARI | Sil. | bARI | ARI | Sil. | bARI |
| Raw | 0.363 | 0.410 | 0.106 | 0.455 | 0.108 | 0.232 | 0.443 | 0.554 | 0.245 | 0.494 | 0.497 | 0.237 |
| Log | 0.406 | 0.535 | 0.179 | 0.400 | 0.151 | 0.113 | 0.429 | 0.686 | 0.243 | 0.460 | 0.524 | 0.161 |
| Total | 0.389 | 0.407 | 0.023 | 0.697 | 0.013 | 0.111 | 0.577 | 0.505 | 0.065 | 0.898 | 0.468 | 0.002 |
| T→L | 0.306 | 0.537 | 0.226 | 0.207 | -0.166 | 0.262 | 0.166 | 0.561 | 0.162 | 0.238 | 0.670 | 0.181 |
| M | 0.467 | 0.418 | 0.094 | 0.908 | 0.383 | -0.019 | 0.719 | 0.619 | 0.060 | 0.805 | 0.490 | 0.051 |
| L→M | 0.398 | 0.529 | 0.195 | 0.344 | 0.109 | 0.191 | 0.444 | 0.621 | 0.218 | 0.311 | 0.668 | 0.358 |
| T→M | 0.481 | 0.419 | 0.080 | 0.799 | 0.296 | 0.052 | 0.719 | 0.619 | 0.060 | 0.805 | 0.490 | 0.051 |
| T→L→M | 0.373 | 0.488 | 0.118 | 0.331 | 0.174 | 0.077 | 0.353 | 0.591 | 0.178 | 0.331 | 0.479 | 0.129 |
| $M^2$ | 0.505 | 0.431 | 0.062 | 0.737 | 0.161 | 0.078 | 0.722 | 0.631 | 0.080 | 0.741 | 0.536 | 0.075 |
| L→ $M^2$ | 0.334 | 0.558 | 0.288 | 0.404 | 0.275 | 0.203 | 0.369 | 0.710 | 0.305 | 0.380 | 0.641 | 0.311 |
| T→ $M^2$ | 0.498 | 0.414 | 0.066 | 0.848 | 0.188 | 0.009 | 0.722 | 0.632 | 0.080 | 0.741 | 0.536 | 0.075 |
| T→L→ $M^2$ | 0.412 | 0.550 | 0.160 | 0.383 | 0.152 | 0.099 | 0.459 | 0.640 | 0.206 | 0.313 | 0.376 | -0.061 |
| l2 | 0.494 | 0.426 | 0.084 | 0.800 | 0.078 | 0.045 | 0.725 | 0.588 | 0.077 | 0.804 | 0.500 | 0.054 |
| L→l2 | 0.355 | 0.568 | 0.261 | 0.390 | 0.360 | 0.194 | 0.371 | 0.719 | 0.336 | 0.409 | 0.514 | 0.274 |
| T→l2 | 0.486 | 0.430 | 0.123 | 0.805 | 0.193 | 0.057 | 0.725 | 0.588 | 0.077 | 0.804 | 0.500 | 0.054 |
| T→L→l2 | 0.413 | 0.550 | 0.165 | 0.380 | 0.083 | 0.100 | 0.460 | 0.650 | 0.206 | 0.422 | 0.375 | 0.109 |
| Z | 0.489 | 0.431 | 0.084 | 0.804 | 0.337 | 0.061 | 0.789 | 0.616 | 0.051 | 0.746 | 0.587 | 0.088 |
| L→Z | 0.322 | 0.554 | 0.249 | 0.347 | 0.297 | 0.232 | 0.358 | 0.726 | 0.320 | 0.375 | 0.664 | 0.318 |
| T→Z | 0.454 | 0.452 | 0.106 | 0.808 | 0.207 | 0.065 | 0.789 | 0.616 | 0.051 | 0.746 | 0.587 | 0.088 |
| T→L→Z | 0.407 | 0.553 | 0.205 | 0.423 | 0.288 | 0.224 | 0.442 | 0.726 | 0.245 | 0.465 | 0.656 | 0.242 |

* R = Raw, L = Log, T = Total, M = Minmax, $M^2$ = Rootmeansquare, l2 = *l*2-norm, Z = Z-score,

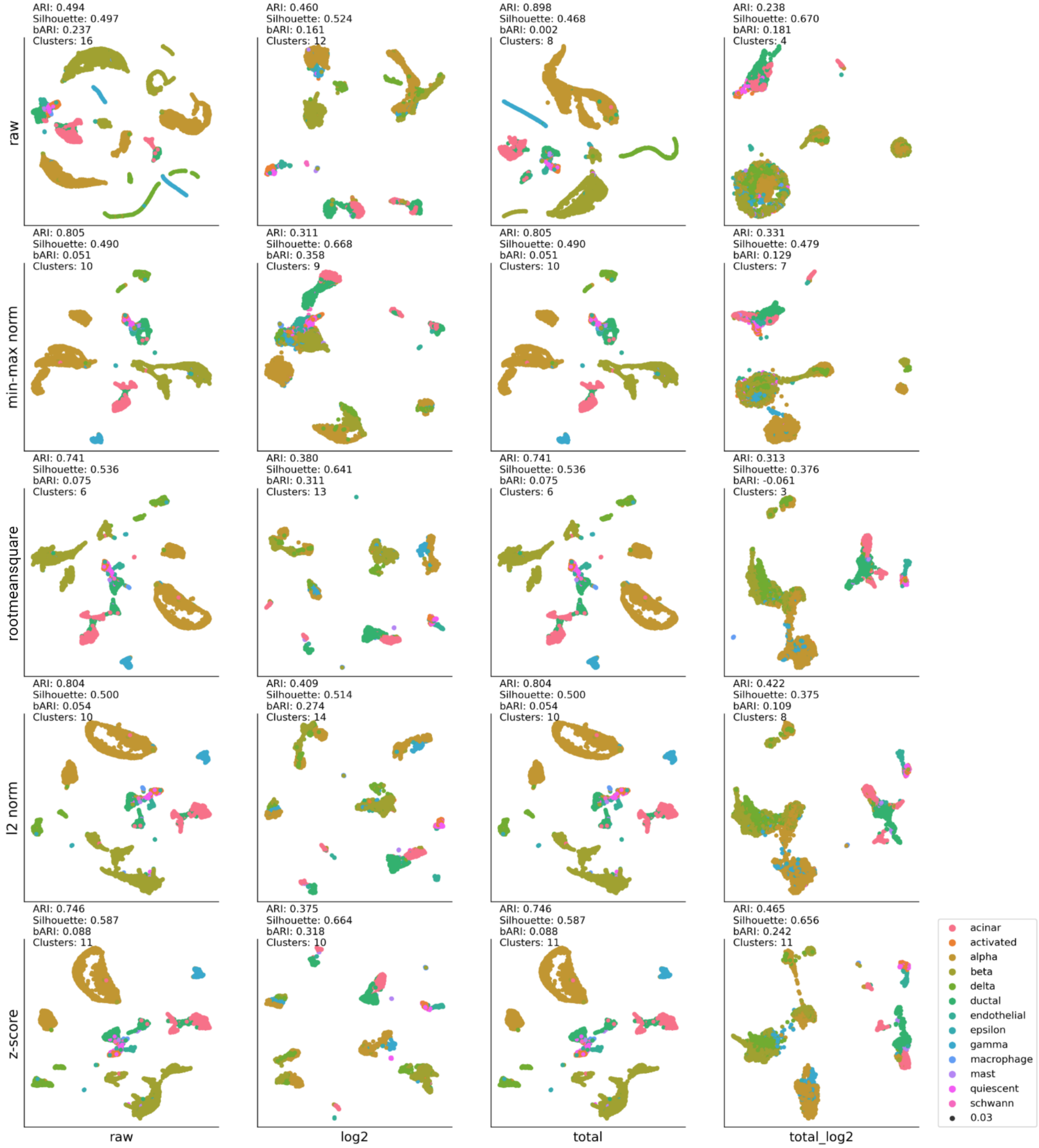


Figure S1: **UMAP representation for comparison of 20 different data transformation.** Five different Human pancreas datasets are integrated and processed for dimensionality reduction and clustering. In this analysis, UMAP and DBSCAN were used. The procedure of data transformation can be read by row to column index order, e.g. result on (2,3) position which has the best ARI (0.914) is preprocessed with Total and after min-max normalization.

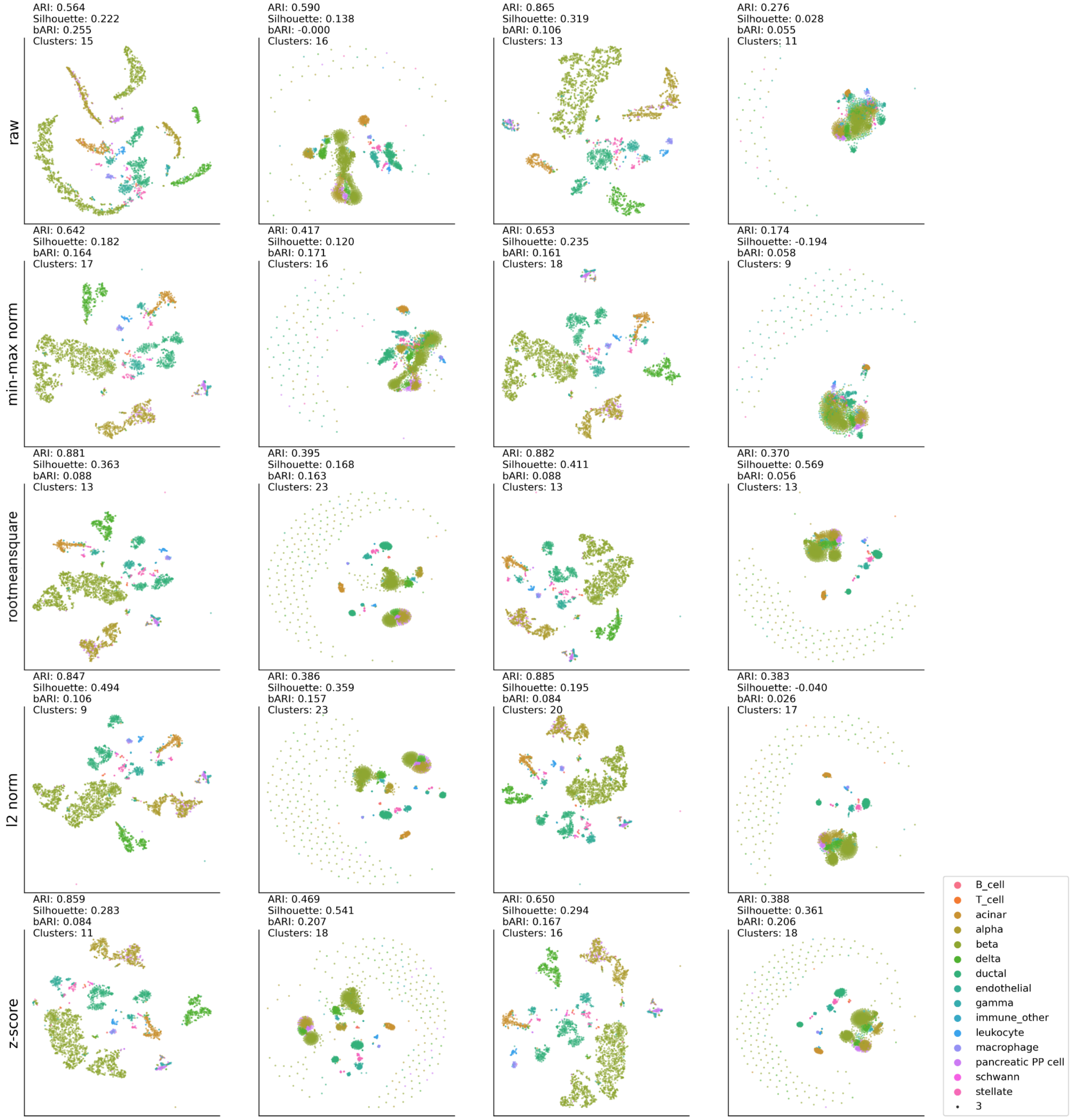
ARI: 0.564
Silhouette: 0.222
bARI: 0.255
Clusters: 15
ARI: 0.590
Silhouette: 0.138
bARI: -0.000
Clusters: 16
ARI: 0.865
Silhouette: 0.319
bARI: 0.106
Clusters: 13
ARI: 0.276
Silhouette: 0.028
bARI: 0.055
Clusters: 11
raw
ARI: 0.642
Silhouette: 0.182
bARI: 0.164
Clusters: 17
ARI: 0.417
Silhouette: 0.120
bARI: 0.171
Clusters: 16
ARI: 0.653
Silhouette: 0.235
bARI: 0.161
Clusters: 18
ARI: 0.174
Silhouette: -0.194
bARI: 0.058
Clusters: 9
min-max norm
ARI: 0.881
Silhouette: 0.363
bARI: 0.088
Clusters: 13
ARI: 0.395
Silhouette: 0.168
bARI: 0.163
Clusters: 23
ARI: 0.882
Silhouette: 0.411
bARI: 0.088
Clusters: 13
ARI: 0.370
Silhouette: 0.569
bARI: 0.056
Clusters: 13
rootmeansquare
ARI: 0.847
Silhouette: 0.494
bARI: 0.106
Clusters: 9
ARI: 0.386
Silhouette: 0.359
bARI: 0.157
Clusters: 23
ARI: 0.885
Silhouette: 0.195
bARI: 0.084
Clusters: 20
ARI: 0.383
Silhouette: -0.040
bARI: 0.026
Clusters: 17
l2 norm
ARI: 0.859
Silhouette: 0.283
bARI: 0.084
Clusters: 11
ARI: 0.469
Silhouette: 0.541
bARI: 0.207
Clusters: 18
ARI: 0.650
Silhouette: 0.294
bARI: 0.167
Clusters: 16
ARI: 0.388
Silhouette: 0.361
bARI: 0.206
Clusters: 18
z-score
B_cell
T_cell
acinar
alpha
beta
delta
ductal
endothelial
gamma
immune_other
leukocyte
macrophage
pancreatic PP cell
schwann
stellate
3

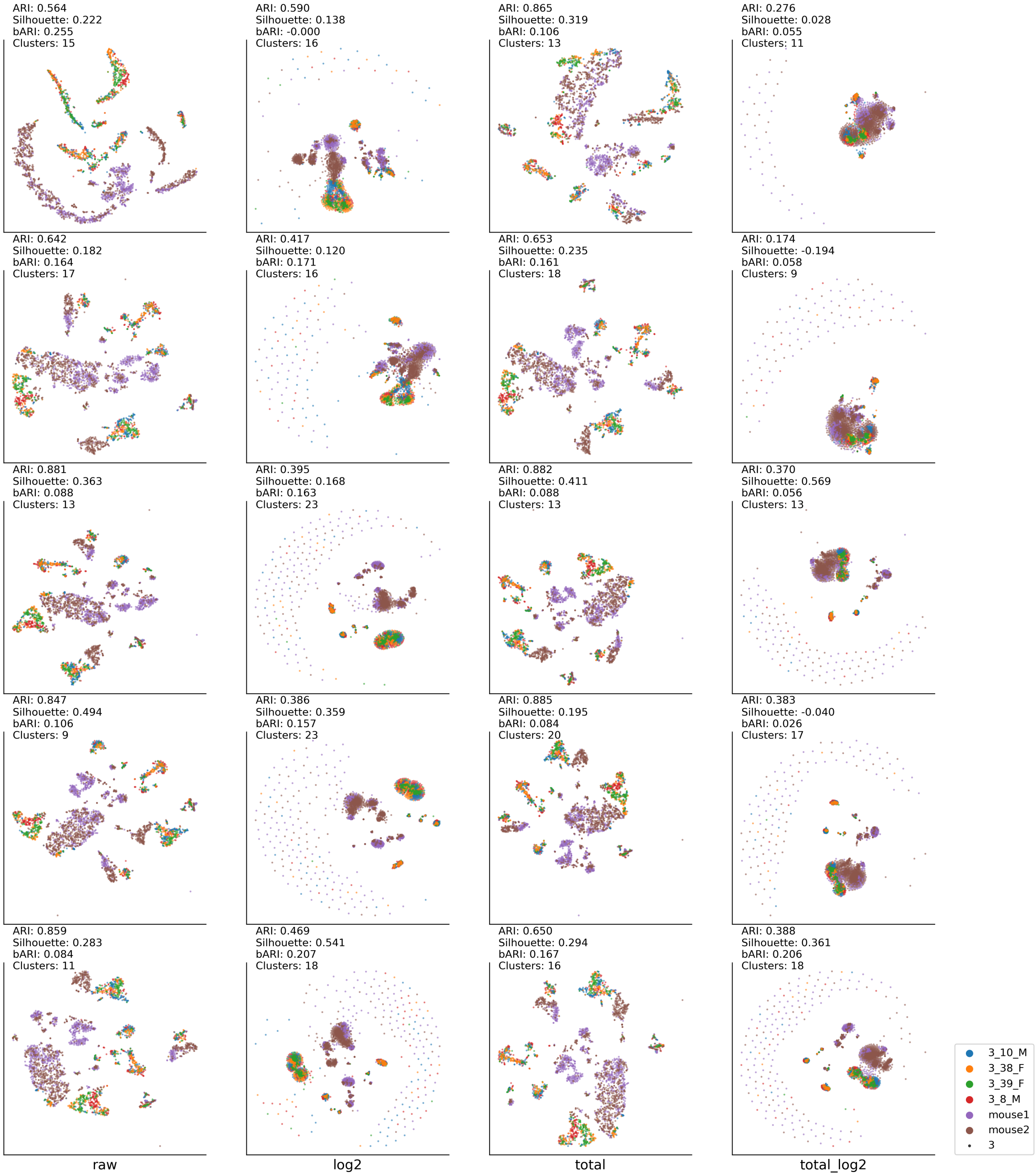


Figure S2: **Mouse Pancreas analysis with t-SNE and DBSCAN** Two mouse pancreas datasets, Tabula Muris pancreas and Baron (Mouse), were integrated. Batch integration was analyzed using t-SNE and DBSCAN. The procedure of data transformation can be read by row to column index order. Batch labels are derived from meta data of original dataset; Tabula Muris: 3-10-M, 3-38-F, 3-39-F, 3-8-M, and Baron (Mouse): mouse1, mouse2.

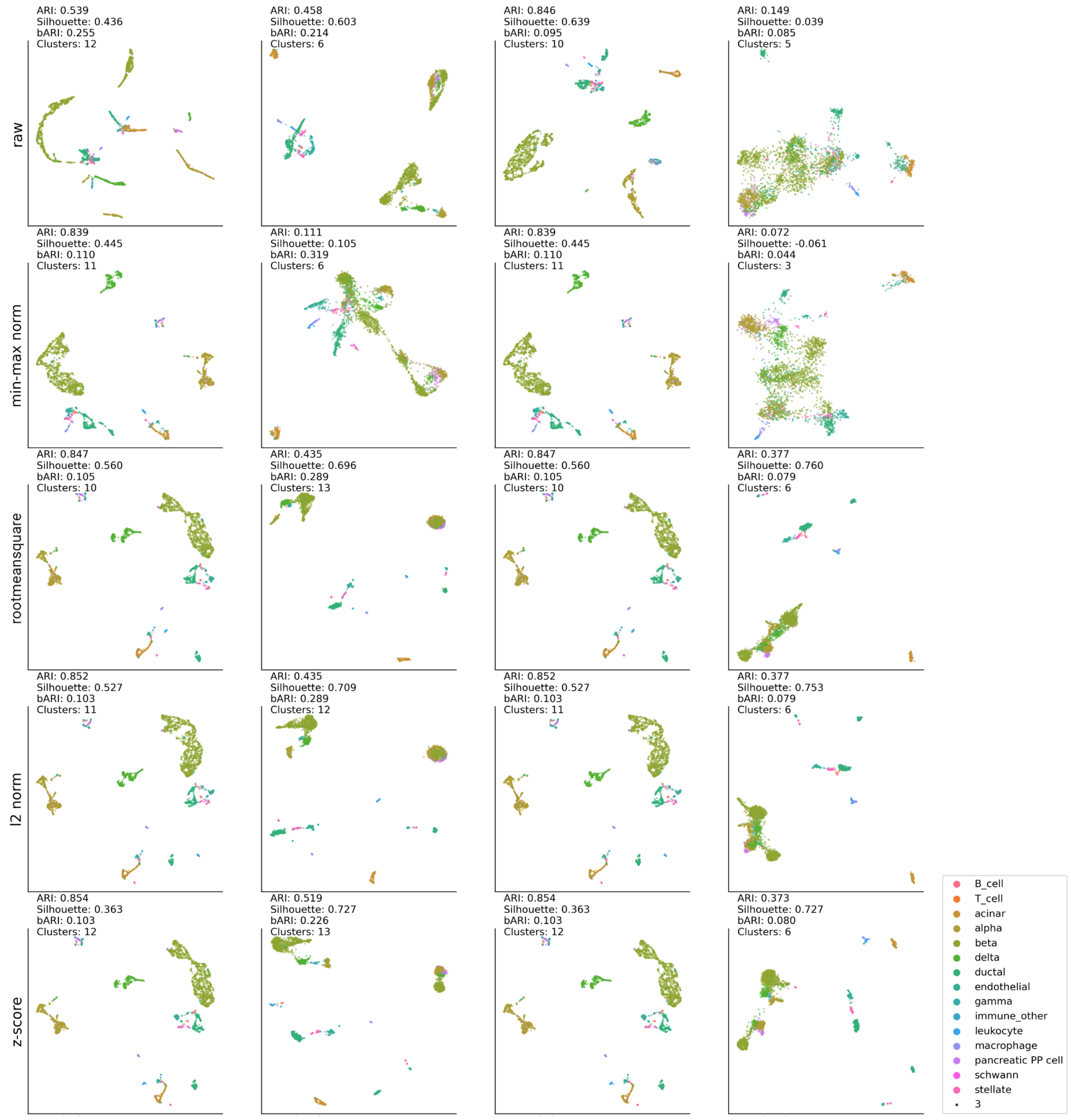
ARI: 0.539
Silhouette: 0.436
bARI: 0.255
Clusters: 12
ARI: 0.458
Silhouette: 0.603
bARI: 0.214
Clusters: 6
ARI: 0.846
Silhouette: 0.639
bARI: 0.095
Clusters: 10
ARI: 0.149
Silhouette: 0.039
bARI: 0.085
Clusters: 5
raw
ARI: 0.839
Silhouette: 0.445
bARI: 0.110
Clusters: 11
ARI: 0.111
Silhouette: 0.105
bARI: 0.319
Clusters: 6
ARI: 0.839
Silhouette: 0.445
bARI: 0.110
Clusters: 11
ARI: 0.072
Silhouette: -0.061
bARI: 0.044
Clusters: 3
min-max norm
ARI: 0.847
Silhouette: 0.560
bARI: 0.105
Clusters: 10
ARI: 0.435
Silhouette: 0.696
bARI: 0.289
Clusters: 13
ARI: 0.847
Silhouette: 0.560
bARI: 0.105
Clusters: 10
ARI: 0.377
Silhouette: 0.760
bARI: 0.079
Clusters: 6
rootmeansquare
ARI: 0.852
Silhouette: 0.527
bARI: 0.103
Clusters: 11
ARI: 0.435
Silhouette: 0.709
bARI: 0.289
Clusters: 12
ARI: 0.852
Silhouette: 0.527
bARI: 0.103
Clusters: 11
ARI: 0.377
Silhouette: 0.753
bARI: 0.079
Clusters: 6
l2 norm
ARI: 0.854
Silhouette: 0.363
bARI: 0.103
Clusters: 12
ARI: 0.519
Silhouette: 0.727
bARI: 0.226
Clusters: 13
ARI: 0.854
Silhouette: 0.363
bARI: 0.103
Clusters: 12
ARI: 0.373
Silhouette: 0.727
bARI: 0.080
Clusters: 6
z-score
B_cell
T_cell
acinar
alpha
beta
delta
ductal
endothelial
gamma
immune_other
leukocyte
macrophage
pancreatic PP cell
schwann
stellate
3

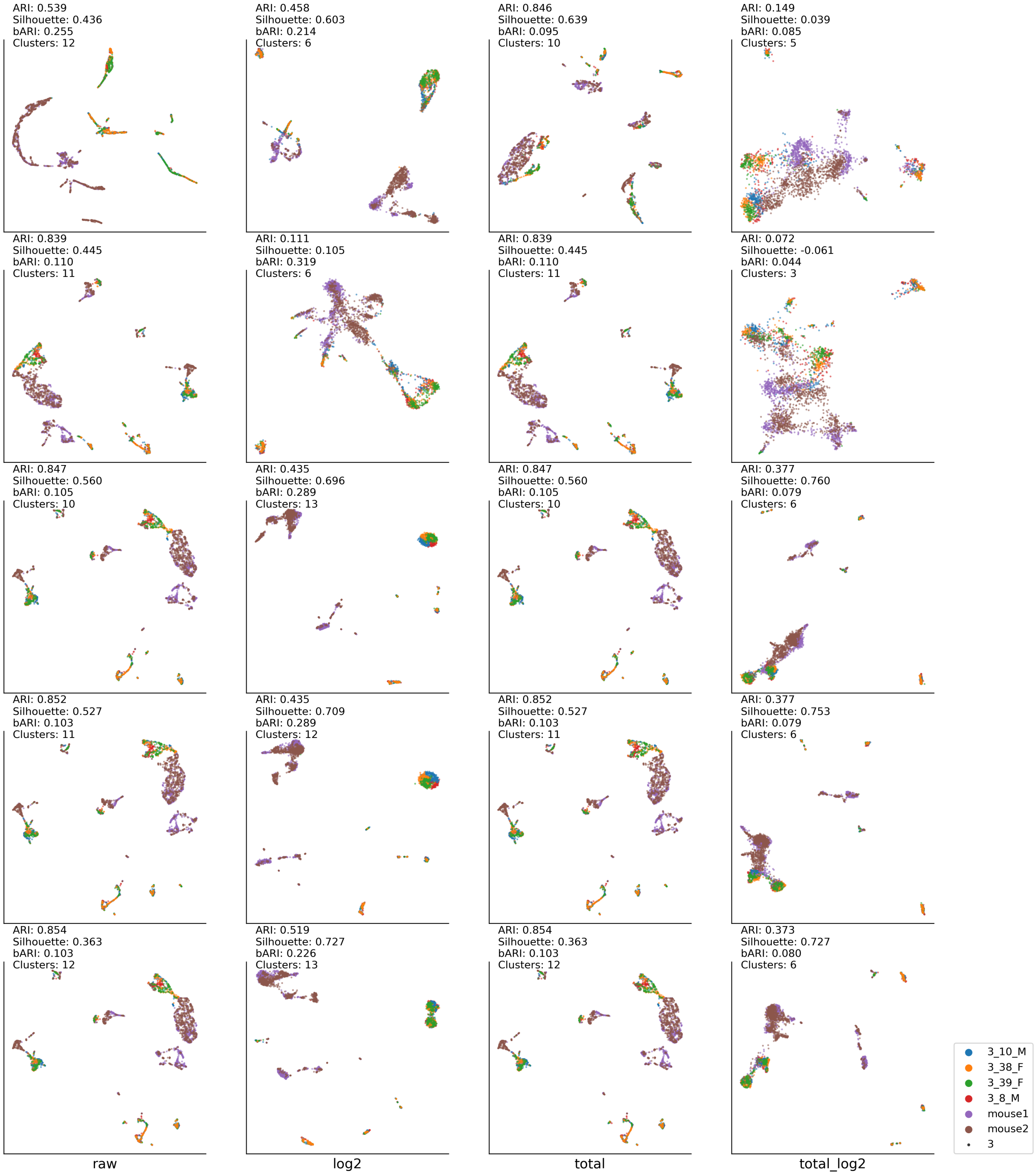


Figure S3: **Mouse Pancreas analysis with UMAP and DBSCAN** Two mouse pancreas datasets, Tabula Muris pancreas and Baron (Mouse), were integrated. Batch integration was analyzed using UMAP and DBSCAN. The procedure of data transformation can be read by row to column index order. Batch labels are derived from meta data of original dataset; Tabula Muris: 3-10-M, 3-38-F, 3-39-F, 3-8-M, Baron (Mouse): mouse1, mouse2.

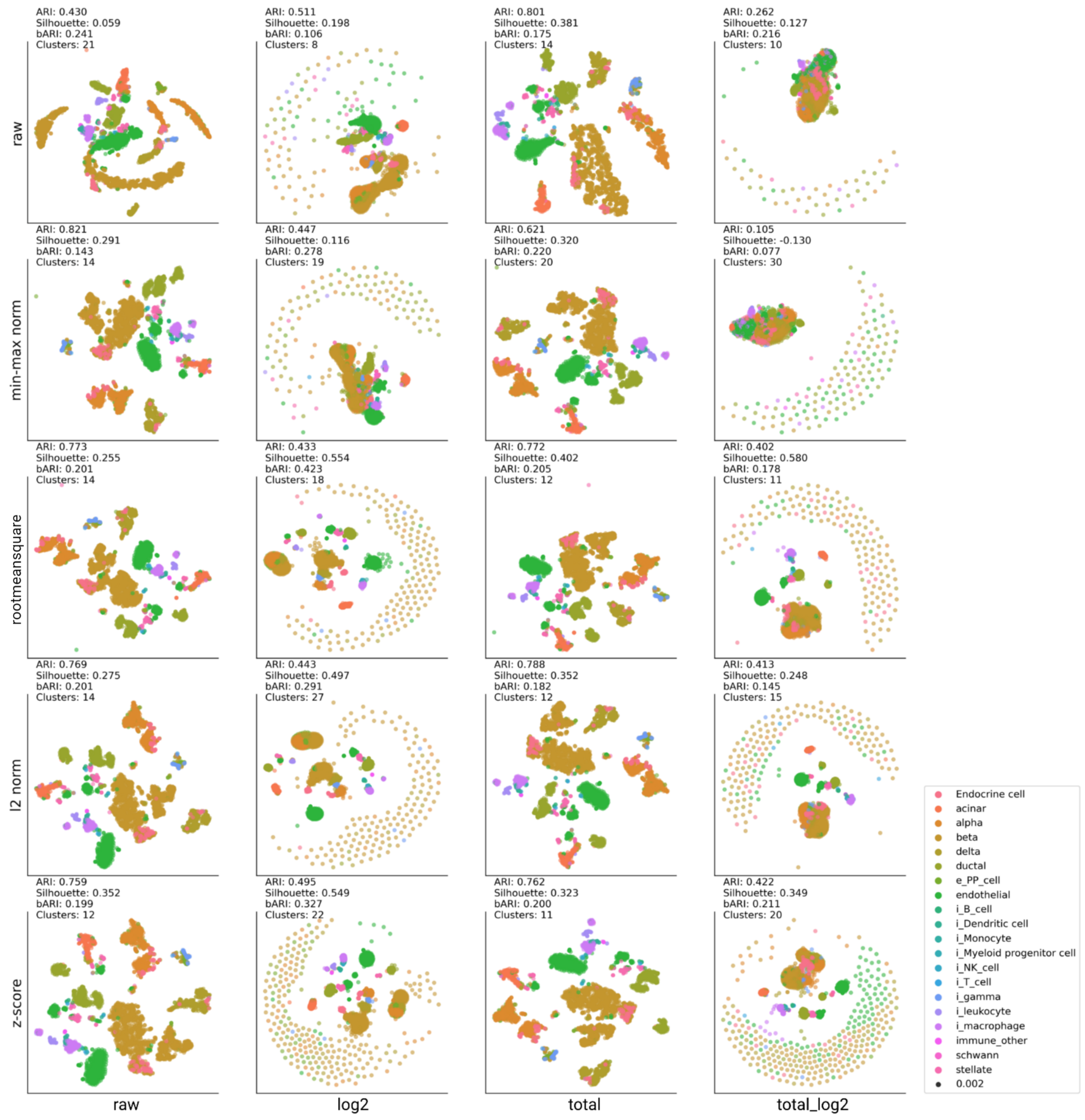
ARI: 0.430
Silhouette: 0.059
bARI: 0.241
Clusters: 21
ARI: 0.511
Silhouette: 0.198
bARI: 0.106
Clusters: 8
ARI: 0.801
Silhouette: 0.381
bARI: 0.175
Clusters: 14
ARI: 0.262
Silhouette: 0.127
bARI: 0.216
Clusters: 10
raw
ARI: 0.821
Silhouette: 0.291
bARI: 0.143
Clusters: 14
ARI: 0.447
Silhouette: 0.116
bARI: 0.278
Clusters: 19
ARI: 0.621
Silhouette: 0.320
bARI: 0.220
Clusters: 20
ARI: 0.105
Silhouette: -0.130
bARI: 0.077
Clusters: 30
min-max norm
ARI: 0.773
Silhouette: 0.255
bARI: 0.201
Clusters: 14
ARI: 0.433
Silhouette: 0.554
bARI: 0.423
Clusters: 18
ARI: 0.772
Silhouette: 0.402
bARI: 0.205
Clusters: 12
ARI: 0.402
Silhouette: 0.580
bARI: 0.178
Clusters: 11
rootmeansquare
ARI: 0.769
Silhouette: 0.275
bARI: 0.201
Clusters: 14
ARI: 0.443
Silhouette: 0.497
bARI: 0.291
Clusters: 27
ARI: 0.788
Silhouette: 0.352
bARI: 0.182
Clusters: 12
ARI: 0.413
Silhouette: 0.248
bARI: 0.145
Clusters: 15
l2 norm
ARI: 0.759
Silhouette: 0.352
bARI: 0.199
Clusters: 12
ARI: 0.495
Silhouette: 0.549
bARI: 0.327
Clusters: 22
ARI: 0.762
Silhouette: 0.323
bARI: 0.200
Clusters: 11
ARI: 0.422
Silhouette: 0.349
bARI: 0.211
Clusters: 20
z-score
raw
log2
total
total_log2
Endocrine cell
acinar
alpha
beta
delta
ductal
e_PP_cell
endothelial
i_B_cell
i_Dendritic cell
i_Monocyte
i_Myeloid progenitor cell
i_NK_cell
i_T_cell
i_gamma
i_leukocyte
i_macrophage
immune_other
schwann
stellate
0.002

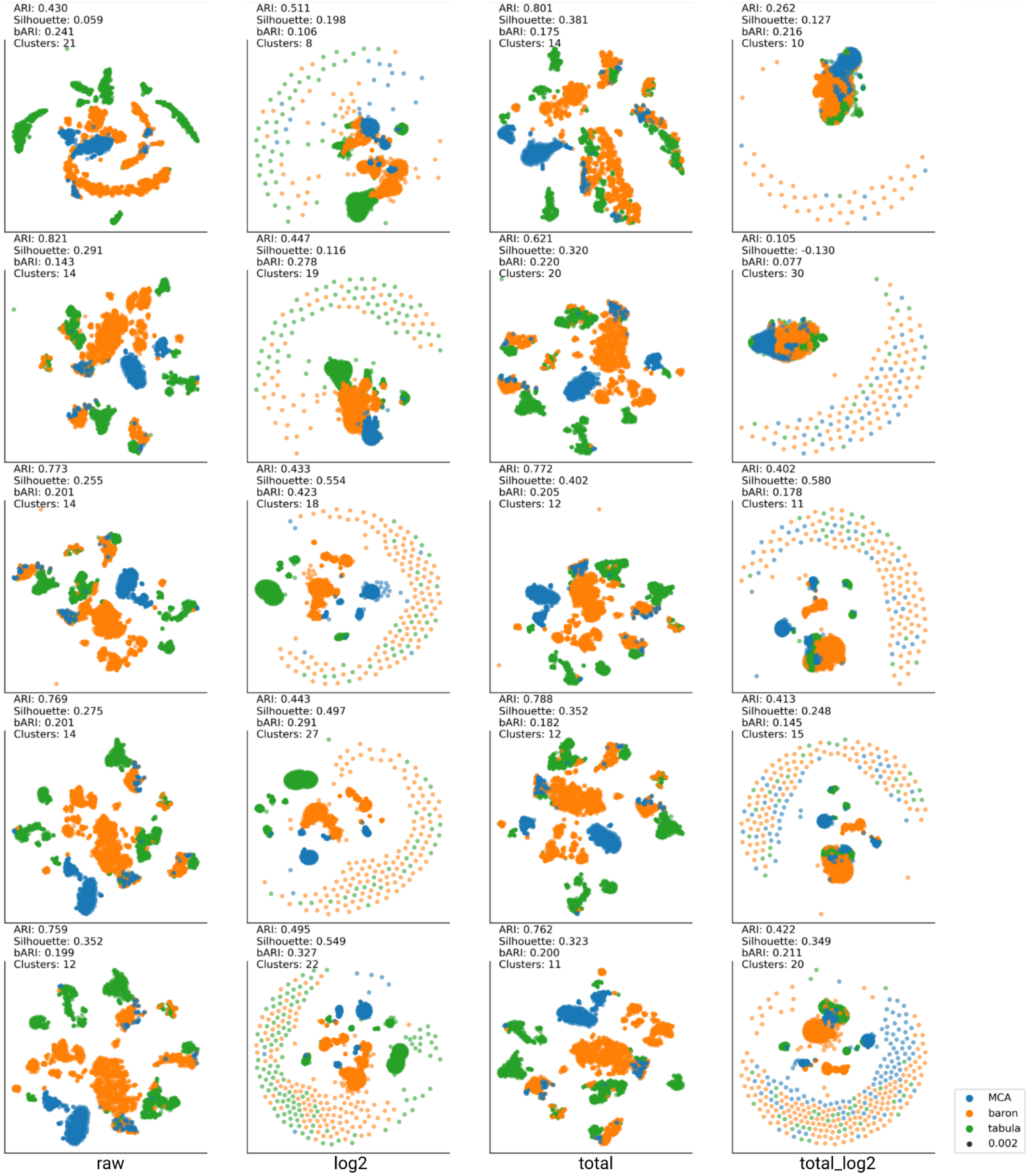


Figure S4: **Three mouse pancreas datasets analysis.** Tabula Muris pancreas, Baron (Mouse), and pancreas of Mouse Cell Atlas dataset were integrated. Batch integrative analysis was done using t-SNE and DBSCAN.

Table S5: **Entire data of integrative analysis with Tabula Muris's two batches.** Two different datasets (FACS and 10x) in Tabula Muris were integrated and analyzed for the impact of data transformation in batch effect. Batch integration and following analysis was done using UMAP and DBSCAN. The best ARI score amongst data transformation methods was marked with green color.

| | Bladder | | | Kideney | | | Limb-musscle | | |
|---|---|---|---|---|---|---|---|---|---|
| | ARI | Sil. | bARI | ARI | Sil. | bARI | ARI | Sil. | bARI |
| Raw | 0.347 | 0.388 | 0.268 | 0.126 | 0.081 | 0.244 | 0.475 | 0.703 | 0.153 |
| Log | 0.747 | 0.688 | 0.004 | 0.472 | 0.775 | 0.025 | 0.746 | 0.649 | 0.074 |
| Total | 0.478 | 0.729 | 0.294 | 0.443 | 0.407 | 0.025 | 0.607 | 0.686 | 0.127 |
| Total→Log | 0.706 | 0.775 | 0.13 | 0.328 | 0.857 | 0.1 | 0.563 | 0.469 | 0.094 |
| Minmax | 0.46 | 0.766 | 0.3 | 0.25 | 0.356 | 0.044 | 0.59 | 0.505 | 0.127 |
| Log→Minmax | 0.812 | 0.734 | -0.002 | 0.791 | 0.707 | 0.026 | 0.834 | 0.72 | 0.027 |
| Total→Minmax | 0.46 | 0.766 | 0.3 | 0.25 | 0.356 | 0.044 | 0.59 | 0.505 | 0.127 |
| Total→Log→Minmax | 0.719 | 0.671 | 0.121 | 0.311 | 0.813 | 0.092 | 0.698 | 0.691 | 0.046 |
| Rootmeansquare | 0.474 | 0.463 | 0.299 | 0.51 | 0.288 | 0.022 | 0.593 | 0.605 | 0.13 |
| Log→Rootmeansquare | 0.78 | 0.703 | -0.001 | 0.481 | 0.8 | 0.025 | 0.844 | 0.652 | 0.025 |
| Total→Rootmeansquare | 0.474 | 0.463 | 0.299 | 0.28 | 0.602 | 0.057 | 0.593 | 0.605 | 0.13 |
| Total→Log→Rootmeansquare | 0.479 | 0.832 | 0.295 | 0.448 | 0.623 | 0.04 | 0.719 | 0.632 | 0.083 |
| *l*2-norm | 0.475 | 0.657 | 0.298 | 0.156 | 0.372 | 0.066 | 0.583 | 0.48 | 0.135 |
| Log→*l*2-norm | 0.748 | 0.783 | 0.003 | 0.48 | 0.798 | 0.025 | 0.844 | 0.674 | 0.025 |
| Total→*l*2-norm | 0.475 | 0.657 | 0.298 | 0.156 | 0.372 | 0.066 | 0.583 | 0.48 | 0.135 |
| Total→Log→*l*2-norm | 0.479 | 0.85 | 0.295 | 0.45 | 0.697 | 0.04 | 0.693 | 0.728 | 0.082 |
| Zscore | 0.474 | 0.622 | 0.299 | 0.474 | 0.391 | 0.016 | 0.599 | 0.664 | 0.131 |
| Log→Zscore | 0.808 | 0.72 | -0.001 | 0.454 | 0.859 | 0.023 | 0.827 | 0.804 | 0.029 |
| Total→Zscore | 0.474 | 0.622 | 0.299 | 0.474 | 0.391 | 0.016 | 0.599 | 0.664 | 0.131 |
| Total→Log→Zscore | 0.479 | 0.852 | 0.295 | 0.448 | 0.586 | 0.039 | 0.693 | 0.718 | 0.082 |

| | Liver | | | Lung | | | Mammary | | |
|---|---|---|---|---|---|---|---|---|---|
| | ARI | Sil. | bARI | ARI | Sil. | bARI | ARI | Sil. | bARI |
| Raw | 0.236 | 0.685 | 0.363 | 0.233 | 0.551 | 0.191 | 0.391 | 0.172 | 0.302 |
| Log | 0.248 | 0.536 | 0.245 | 0.435 | 0.48 | 0.067 | 0.7 | 0.792 | 0.116 |
| Total | 0.392 | 0.53 | 0.14 | 0.365 | 0.452 | -0.055 | 0.696 | 0.732 | 0.175 |
| Total→Log | 0.641 | 0.697 | 0.128 | 0.169 | 0.57 | 0.252 | 0.597 | 0.734 | 0.269 |
| Minmax | 0.234 | 0.577 | 0.362 | 0.31 | 0.43 | 0.176 | 0.521 | 0.581 | 0.275 |
| Log→Minmax | 0.82 | 0.648 | 0.084 | 0.744 | 0.698 | 0.011 | 0.884 | 0.71 | 0.104 |
| Total→Minmax | 0.234 | 0.577 | 0.362 | 0.31 | 0.43 | 0.176 | 0.521 | 0.581 | 0.275 |
| Total→Log→Minmax | 0.428 | 0.47 | 0.107 | 0.137 | -0.032 | -0.061 | 0.621 | 0.835 | 0.264 |
| Rootmeansquare | 0.233 | 0.605 | 0.363 | 0.362 | 0.586 | 0.148 | 0.686 | 0.74 | 0.184 |
| Log→Rootmeansquare | 0.656 | 0.495 | 0.022 | 0.752 | 0.729 | 0.012 | 0.897 | 0.807 | 0.094 |
| Total→Rootmeansquare | 0.233 | 0.605 | 0.363 | 0.362 | 0.586 | 0.148 | 0.686 | 0.74 | 0.184 |
| Total→Log→Rootmeansquare | 0.869 | 0.757 | 0.063 | 0.636 | 0.704 | 0.062 | 0.619 | 0.841 | 0.236 |
| *l*2-norm | 0.234 | 0.564 | 0.363 | 0.398 | 0.679 | 0.139 | 0.672 | 0.711 | 0.184 |
| Log→*l*2-norm | 0.656 | 0.559 | 0.022 | 0.772 | 0.572 | 0.016 | 0.896 | 0.818 | 0.094 |
| Total→*l*2-norm | 0.234 | 0.564 | 0.363 | 0.398 | 0.679 | 0.139 | 0.672 | 0.711 | 0.184 |
| Total→Log→*l*2-norm | 0.878 | 0.724 | 0.065 | 0.645 | 0.725 | 0.064 | 0.619 | 0.84 | 0.237 |
| Zscore | 0.235 | 0.556 | 0.363 | 0.368 | 0.614 | 0.147 | 0.719 | 0.683 | 0.189 |
| Log→Zscore | 0.867 | 0.46 | 0.065 | 0.76 | 0.753 | 0.018 | 0.895 | 0.835 | 0.095 |
| Total→Zscore | 0.235 | 0.556 | 0.363 | 0.368 | 0.614 | 0.147 | 0.687 | 0.777 | 0.185 |
| Total→Log→Zscore | 0.876 | 0.515 | 0.064 | 0.477 | 0.616 | 0.124 | 0.642 | 0.85 | 0.255 |

| | Marrow | | | Spleen | | | Thymus | | |
|---|---|---|---|---|---|---|---|---|---|
| | ARI | Sil. | bARI | ARI | Sil. | bARI | ARI | Sil. | bARI |
| Raw | 0.211 | 0.232 | 0.161 | 0.344 | 0.623 | 0.285 | 0.007 | 0.539 | 0.344 |
| Log | 0.246 | 0.363 | -0.059 | 0.67 | 0.179 | -0.057 | NA | NA | NA |
| Total | 0.292 | 0.405 | 0.145 | 0.417 | 0.35 | 0.273 | 0.034 | 0.466 | 0.422 |
| Total→Log | 0.259 | 0.488 | 0.129 | 0.058 | 0.814 | 0.41 | NA | NA | NA |
| Minmax | 0.228 | 0.432 | 0.214 | 0.06 | 0.659 | 0.429 | NA | NA | NA |
| Log→Minmax | 0.041 | 0.174 | -0.036 | 0.921 | 0.571 | -0.008 | NA | NA | NA |
| Total→Minmax | 0.228 | 0.432 | 0.214 | 0.06 | 0.659 | 0.429 | NA | NA | NA |
| Total→Log→Minmax | 0.082 | 0.041 | 0.318 | 0 | 0 | 10 | 0.002 | 0.362 | 0.633 |
| Rootmeansquare | 0.316 | 0.463 | 0.15 | 0.489 | 0.475 | 0.298 | NA | NA | NA |
| Log→Rootmeansquare | 0.243 | 0.458 | -0.055 | 0.233 | 0.323 | -0.009 | 0.117 | 0.126 | -0.004 |
| Total→Rootmeansquare | 0.316 | 0.463 | 0.15 | 0.489 | 0.475 | 0.298 | NA | NA | NA |
| Total→Log→Rootmeansquare | 0.197 | 0.265 | 0.234 | 0.217 | 0.274 | -0.009 | 0.536 | -0.235 | 0.004 |
| *l*2-norm | 0.316 | 0.513 | 0.145 | 0.482 | 0.486 | 0.301 | NA | NA | NA |
| Log→*l*2-norm | 0.238 | 0.386 | -0.057 | 0.234 | -0.07 | -0.009 | 0.117 | 0.172 | -0.004 |
| Total→*l*2-norm | 0.316 | 0.513 | 0.145 | 0.482 | 0.486 | 0.301 | NA | NA | NA |
| Total→Log→*l*2-norm | 0.193 | 0.245 | 0.232 | 0.218 | 0.276 | -0.009 | 0.545 | 0.054 | 0.003 |
| Zscore | 0.315 | 0.486 | 0.145 | 0.468 | 0.328 | 0.297 | NA | NA | NA |
| Log→Zscore | 0.262 | 0.158 | -0.045 | 0.947 | 0.648 | -0.005 | NA | NA | NA |
| Total→Zscore | 0.315 | 0.503 | 0.145 | 0.489 | 0.43 | 0.298 | NA | NA | NA |
| Total→Log→Zscore | 0.198 | 0.528 | 0.234 | 0.223 | 0.139 | -0.008 | 0.532 | -0.219 | 0.004 |

| | Tongue | | | Trachya | | |
|---|---|---|---|---|---|---|
| | ARI | Sil. | bARI | ARI | Sil. | bARI |
| Raw | 0.182 | -0.023 | 0.229 | 0.112 | 0.072 | 0.091 |
| Log | 0.231 | -0.123 | -0.024 | 0.062 | 0.084 | -0.004 |
| Total | 0.219 | -0.008 | 0.218 | 0.51 | 0.167 | 0.093 |
| Total→Log | 0.076 | -0.132 | 0.046 | 0.683 | 0.656 | 0.01 |
| Minmax | 0.221 | 0.002 | 0.223 | 0.207 | 0.642 | 0.114 |
| Log→Minmax | 0.183 | 0.048 | 0.076 | 0.077 | -0.082 | -0.004 |
| Total→Minmax | 0.167 | -0.128 | 0.189 | 0.207 | 0.642 | 0.114 |
| Total→Log→Minmax | 0.083 | -0.174 | 0.076 | 0.735 | 0.673 | 0.003 |
| Rootmeansquare | 0.241 | 0.055 | 0.166 | 0.398 | 0.488 | 0.081 |
| Log→Rootmeansquare | 0.168 | 0.136 | 0.141 | 0.749 | 0.208 | 0.088 |
| Total→Rootmeansquare | 0.292 | 0.175 | 0.206 | 0.398 | 0.488 | 0.081 |
| Total→Log→Rootmeansquare | 0.092 | -0.023 | 0.251 | 0.743 | 0.352 | 0.085 |
| *l*2-norm | 0.177 | -0.05 | 0.188 | 0.209 | 0.586 | 0.116 |
| Log→*l*2-norm | 0.201 | -0.039 | -0.024 | 0.747 | 0.254 | 0.089 |
| Total→*l*2-norm | 0.264 | 0.188 | 0.174 | 0.209 | 0.586 | 0.116 |
| Total→Log→*l*2-norm | 0.088 | -0.155 | 0.047 | 0.742 | 0.508 | 0.085 |
| Zscore | 0.178 | 0.017 | 0.188 | 0.401 | 0.637 | 0.082 |
| Log→Zscore | 0.205 | -0.227 | -0.011 | 0.73 | 0.564 | 0.09 |
| Total→Zscore | 0.241 | 0.071 | 0.166 | 0.401 | 0.637 | 0.082 |
| Total→Log→Zscore | 0.092 | 0.021 | 0.248 | 0.743 | 0.71 | 0.084 |

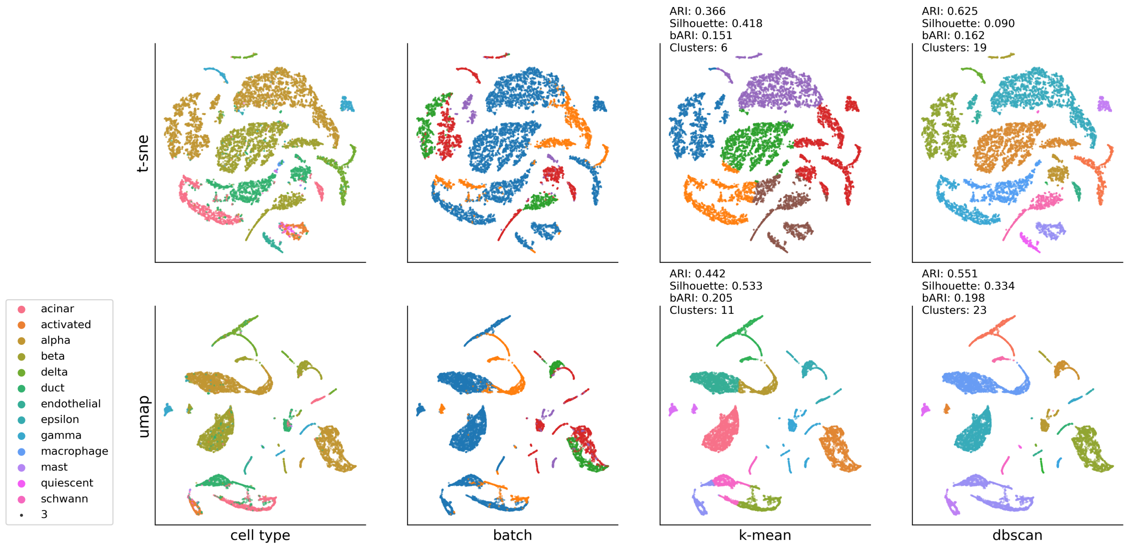


Figure S5: **Latent representation using Autoencoder without data transformation.** The model was trained by MAML 200 episodes with the Baron dataset and evaluated on HP datasets. Human pancreas datasets were not processed. Latent vector size is set to 128.

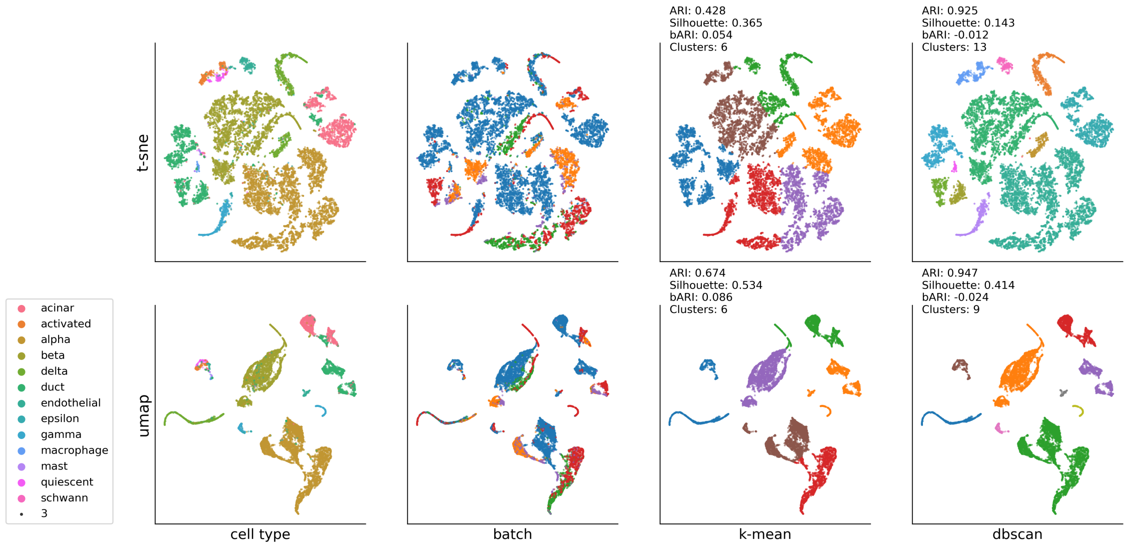


Figure S6: **Latent representation using Autoencoder with Total data transformation.** The model was trained by MAML 200 episodes with the Baron dataset and evaluated on the HP datasets. Human pancreas datasets were transformed with Total. Latent vector size is set to 128.

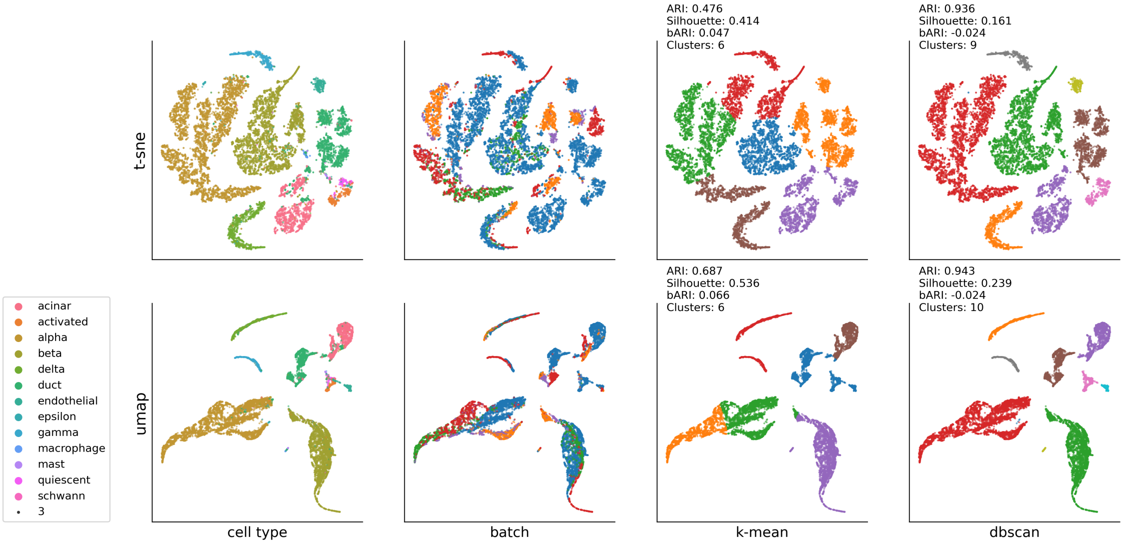


Figure S7: **Latent representation using Variational Autoencoder with Total data transformation.** The model was trained by MAML 200 episodes with the Baron dataset and evaluated on the HP datasets. Human pancreas dataset is transformed with Total method. Latent vector size is set to 128.

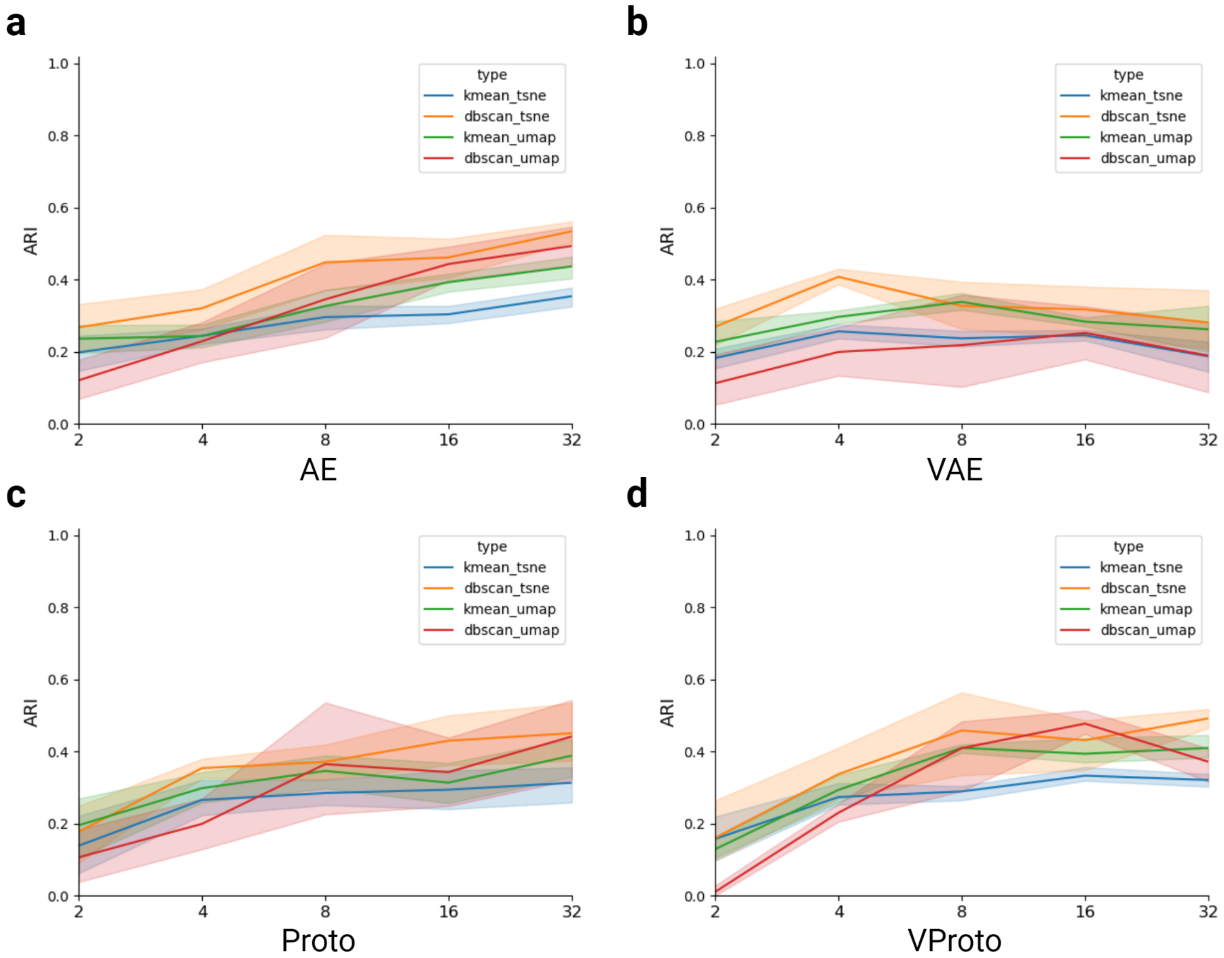


Figure S8: **The performance of deep neural networks models with different size of latent vectors.** When RAW expression count is used, the AE and VAE models showed relatively poor ARI compared to Total transformed data (Figure 4). All models were trained with 200 episodes of MAML training with Baron datasets and evaluated.
